# *de Novo* Sequencing of Antibodies for Identification of Neutralizing Antibodies in Human Plasma Post SARS-CoV-2 Vaccination

**DOI:** 10.1101/2024.03.14.583523

**Authors:** Thierry Le Bihan, Teressa Nunez de Villavicencio Diaz, Chelsea Reitzel, Victoria Lange, Minyoung Park, Emma Beadle, Lin Wu, Marko Jovic, Rosalin M. Dubois, Amber L. Couzens, Jin Duan, Xiaobing Han, Qixin Liu, Bin Ma

## Abstract

We present a method for sequencing polyclonal IgG enriched from human plasma, employing a combination of *de novo* sequencing, proteomics, bioinformatics, protein separation, sequencing, and peptide separations. Our study analyzes a single patient’s IgG antibody response triggered by the Moderna Spikevax mRNA COVID-19 vaccine. From the sequencing data of the natural polyclonal response to vaccination, we generated 12 recombinant antibodies. Six derived recombinant antibodies, including four generated with *de novo* sequencing, exhibited similar or higher binding affinities than the original natural polyclonal antibody. Our neutralization tests revealed that the six antibodies possess neutralizing capabilities against the target antigen. This research provides insights into sequencing polyclonal IgG antibodies while highlighting the effectiveness and potential of our approach in generating recombinant antibodies with robust binding affinity and neutralization capabilities. Our proposed approach is an advancement in characterizing the IgG response by directly investigating the circulating pool of IgG without relying exclusively on the B-cell repertoire or population. This is crucial as the B-cell analysis may not accurately represent the circulating antibodies. Interestingly, a large proportion (80 to 90%) of the human antibody sequences generated against SARS-CoV-2 in the literature have been derived solely from B-cell analysis. Therefore, the ability to offer a different perspective is crucial in gaining a comprehensive understanding of the IgG response.

**Significance Statement:** We investigate human IgG targeting the receptor binding domain using *de novo* proteomics. The peripheral B-cell repertoire may not adequately cover all the circulating IgG for human IgG sequencing. Our approach overcomes this limitation by using a *de novo* protein sequencing on top of standard proteomics. We obtained distinct *de novo* sequences, showcasing our method’s potential. The recombinant proteins we generate possess traits comparable to or surpassing the naturally occurring polyclonal antibodies (pAbs). This study highlights similarities and differences between IgG populations in blood and circulating B-cells, which is crucial for future biologics development.

## Introduction

Antibodies are highly selective molecules and attractive for biotherapeutics and diagnostic assays. Therapeutic antibodies have been used to treat cancer, infections, and autoimmune diseases (1). Polyclonal antibodies (pAbs) are usually produced with immunized animals, which suffer from the batch-to-batch variability problem (2). Monoclonal antibodies (mAbs) overcome the variability problem and have traditionally been produced by hybridoma (3). More recently, mAbs have been developed through phage display (4, 5). There is a growing interest in discovering antibodies by interrogating the natural immune response of animals or humans using ex vivo single B-cell sequencing (4, 6) and the combined use of B-cell sequencing with proteomics (7–10). This manuscript explores the possibility of *de novo* sequencing mAbs directly from a pAb mixture as a complementary approach to overcome existing technological limitations.

Phage display technology uses bacteriophages to incorporate genetic sequences encoding antibodies, allowing for the display of antibody variants on the surface of the phage, enabling antibody discovery and affinity maturation. The application of phage display has significantly influenced antibody engineering, particularly following the approval of adalimumab. *In vitro* display technologies offer broad immune libraries for antibody production, but unnatural heavy and light chain pairing poses challenges in developability and affinity. Around 18% of FDA-licensed monoclonal antibodies have been developed through phage display(11).

More therapeutic antibodies have been discovered by leveraging the B-cell repertoire of the natural immune response in transgenic animals or, more effectively, in human patients. For instance, 19 out of the 28 FDA-approved human monoclonal antibodies from 2002 to 2018 were derived from “humanized” transgenic mice (1). However, these mouse-derived fully human antibodies are not completely non-immunogenic in humans (12). Antibodies generated through natural human immune responses undergo tolerization in humans, rendering them safer and more effective than those derived from other species (4). The SARS-CoV-2 pandemic has led to the isolation of B-cells from convalescent patients, resulting in the discovery of antibodies like Tixagevimab-Cilgavimab, Bamlanivimab, and Bebtelovimab. Furthermore, Ofatumumab, Daratumumab, and Ustekinumab are human B-cell-derived antibodies targeting CD20, CD38, and Il-12/23, respectively (13, 14). Additionally, VRC01, a broadly neutralizing HIV antibody, was isolated from HIV-positive B-cells (15).

While B-cell sequencing has been effective for antibody discovery, peripheral B-cells may not capture the full diversity of peripheral antibodies. Most Long-Lived Plasma B-cells (LLPCs) reside in the bone marrow and lymphoid organs, producing high-affinity antibodies for an extended period. Obtaining LLPCs from tissues can be challenging. Peripheral blood, with a small fraction of B-cells, provides limited information. Peripheral blood, with a small fraction of B-cells, provides limited information. Studies estimate that 2% (16) to none (17) matches the circulating IgG. Not all circulating B-cells produce antibodies (18). Overlap between circulating IgG and peripheral blood B-cells varies based on factors like antigen, immune status, or timing. Relying solely on blood-circulating B-cells for development candidates is challenging, unreliable, and potentially random. B-cell sequencing offers insights into the immune system, but correlations and limitations must be considered for immune repertoire characterization. Ultimately, immune protection depends on serum antibodies, not B-cell receptors.

B-cells from various organs produce polyclonal antibodies (pAbs). IgGs, with a long half-life, are highly abundant in blood circulation, facilitating sampling. Traditional affinity purification on a solid substrate enriches antibodies to a specific antigen. High-affinity polyclonal antibodies (pAbs) can then be extracted using this technique, which reduces the complexity of the pAbs that can then be analyzed using mass spectrometry (MS). Matching the MS data with the sequence database produced by B-cell repertoire sequencing helps discover antigen-specific antibodies. This proteogenomics approach has generated monoclonal antibodies from animals and humans (7–9). The proteogenomics approach improves upon B-cell approaches by identifying antibodies in the serum. However, it cannot recover circulating antibodies absent from the B-cell sequencing database, potentially comprising a significant quantity. This manuscript’s results will demonstrate the existence of these uncovered antibodies.

One potential solution to the problem is *de novo* sequencing of antibody proteins, without relying on a reference database. *De novo* sequencing of monoclonal antibodies has been extensively investigated and studied (19–21) and has reached sufficient maturity for commercial services. Typically, these methods involve enzymatic digestion of the mAb to generate overlapping peptides, *de novo* sequencing of each peptide’s tandem mass spectra (MS/MS), and assembly of the protein sequence by overlapping the peptide sequences.

However, due to unresolved challenges, *de novo* sequencing studies of polyclonal antibodies have been limited. Previous efforts have not successfully discovered functional antibodies matching the affinity of the original pAb. Guthals et al. (17) attempted pAb *de novo* sequencing using a procedure similar to mAb sequencing, but the resulting monoclonal antibodies had significantly lower affinity, thus potentially incorrect sequences. Additionally, their starting pAb mixture was more oligoclonal than polyclonal, limiting the method’s applicability to complex pAbs. Bondt et al. (22) recently used a similar approach, including middle-down MS, but did not provide data demonstrating affinity or neutralization effectiveness of the resulting antibodies. These earlier studies were ambitious but revealed the difficulty of pAb *de novo* sequencing. Our experience suggests that assembling the CDRs into chains and pairing the heavy and light chains accurately poses one of the main challenges. The large number of mAb clones in a pAb leads to a significant increase in combinations, which earlier methods fail to address. Further discussion of the challenges and limitations of existing approaches can be found in the manuscript’s discussion section.

The present study sequences a COVID-19-vaccinated patient’s polyclonal antibodies using multiple methods. One of the approaches is *de novo* sequencing, which combines the mass spectrometry data generated in several orthogonal experiments to address the aforementioned difficulties. These include the bottom-up proteomics to provide accurate sequence of local regions; peptide chemistry (i.e., amino-ethylation of cysteines and peptide C-terminal labeling with methyl-arginine) to maximize sequence coverage and to enhance the w-ions to distinguish Ile and Leu; middle-down proteomics to confirm longer stretches of sequences; and protein separation and label-free quantitation, as well as digestion under non-reductive conditions, all to increase confidence in assembling full-length heavy and light chains. Additionally, we propose a strategy for heavy-light chain pairing. In addition to the *de novo* sequencing approach, we use a B-cell repertoire analysis to evaluate its overlap with the proteomics dataset. Finally, we expressed the obtained antibody sequences and confirmed their binding affinity and neutralization of the Fluc-GFP pseudovirus.

## Results

Different processing methods were used on proteomics data (Fig. 1A). The data was initially analyzed conventionally and compared to the IgSeq/B-cell database for the same individual. In addition, a *de novo* analysis was performed on the proteomics dataset. Some recombinant antibodies were generated from this analysis and subsequently tested for antigen binding and neutralization using ELISA, SPR, and pseudovirus neutralization assays. The sequencing protocol for *de novo* analysis is outlined in Fig. 1B. Multiple proteases were used to generate overlapping peptides from antibody fractions enriched with antigens. Short reads were then used to target the heavy and light chains’ CDR1, CDR2, and CDR3 regions. We extended the short-read assemblies based on high peptide overlap confidence and stopped when ambiguities were encountered. The accuracy of longer sequences was verified using a customized FASTA database. Furthermore, this database confirmed the assembly of multiple CDRs in chains through middle-down analysis. Non-reducing experiments were performed to identify sequence pairs for CDR1 and CDR3. Transitioning from short-read to longer-read assembly was facilitated by native gel electrophoresis of the polyclonal antibody, aiding in pairing sequence-specific peptides for the heavy and light chains.

**Figure 1:**
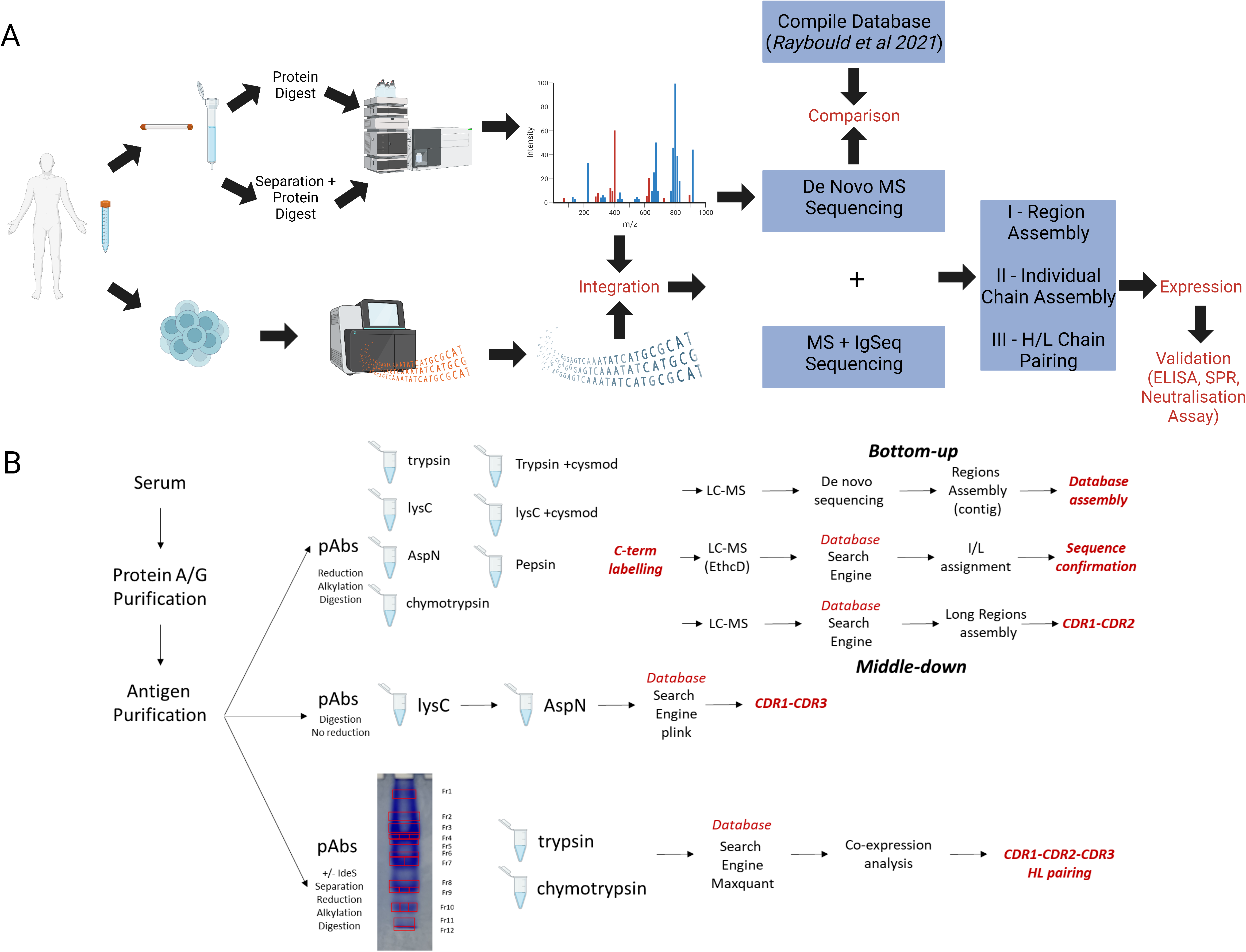
Schematic overview of the sequencing pipeline. **Fig. 1A** depicts the overview analysis of human antibody repertoires from antigen-specific IgG and peripheral blood. Two different types of samples were taken and analyzed; the bottom branch is the procedure of generating a reference database of immunoglobulin V-region by next-generation sequencing (NGS) of the immunized individual’s B-cell repertoire. The branch above describes the analysis of soluble serum IgG. This comparison and functional characterization of the two antibody repertoires provide a different perspective on studying humoral response. This process involves the purification and the proteomic analysis of affinity-purified serum antibody (*de novo*, top) alongside VH: VL pairing and NGS of peripheral B cell V gene repertoires (BCR-seq, bottom of **Fig. 1A**). sThe proteomics dataset was analyzed using the B-Cell repertoire. A *de novo* sequencing analysis of the proteomics dataset was performed in parallel. The two datasets were compared to the Compile database from Raybould et al. 2021 (CoV-AbDab), comprising a comprehensive list of antibody sequences developed in the COVID context. The process of assembling antibodies can be divided into three steps: (1) Different CDR regions assembled, (2) Intact individual chain assembly performed, and (3) heavy and light chain pairing. The generated sequences were expressed recombinantly, and their performance was evaluated. **Fig. 1B** shows a breakdown of the different proteomics experiments performed under the *de novo* sequencing.

### RNA Sequencing

This study screened three consenting SARS-CoV-2 vaccinees (patient information in *SI Appendix table S1*). Screening targeted RBD protein-specific IgG antibodies. Each patient provided 10 ml of plasma for analysis and 8 ml of blood for B-Cell repertoire and IgSeq. A secondary goat anti-human IgG Fc ELISA was used to detect human IgG antibodies. Patient “522” showed the strongest response, followed by “422” and “922”. Our study focused on patient 522 (Fig. 2A). The TakaraBio SMARTer Human BCR IgG IgM H/K/L Profiling Kit was used to create NGS libraries. RNA from patient “522”’s PBMC was used for library construction. Immunoglobulin heavy and light chain variable regions were generated using the method described in the Materials and Methods section. Sequences for the heavy, kappa, and lambda chains were used for proteomics analysis. We obtained 361,895 unique heavy-chain sequences. Kappa and lambda chains yielded 186,247 and 237,074 unique sequences, respectively. Duplicate acquisitions were made for the heavy chain to ensure reproducibility.

**Figure 2:**
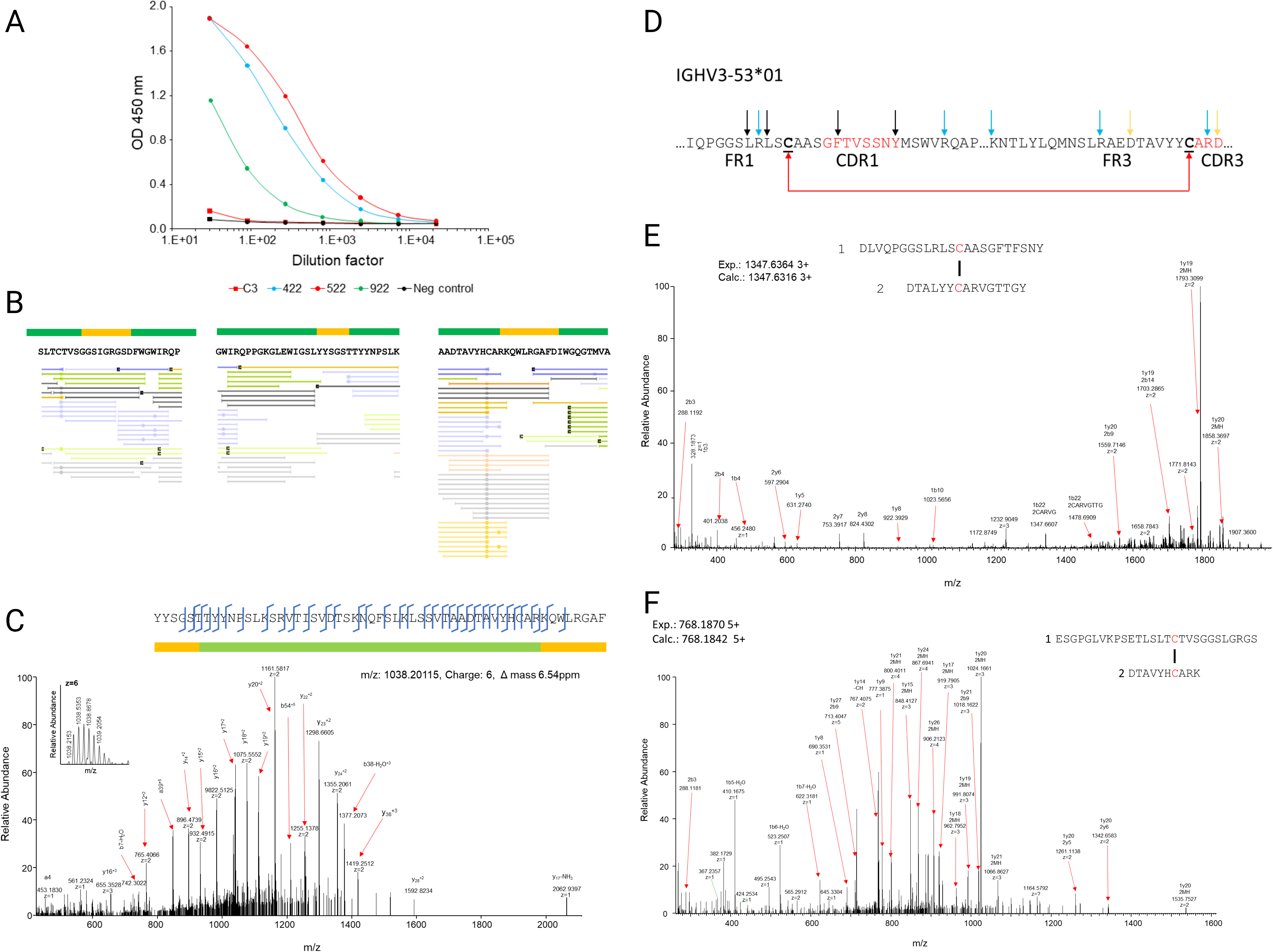
Sample assessment and the first steps of *de novo* sequencing. **Fig. 2A** consists of a plasma titration for antibodies against the receptor binding domain, RBD of Sars-Cov2. Three patient samples (422, 522, and 922) were benchmarked against an internal in-house standard (C3) and a negative control. **Fig. 2B** Illustrates the construct of the *de novo* assembly of “contig” around different CDRs (CDR1 left, CDR2 middle, and CDR3 right). The green region is the framework region, while the orange one is the CDRs for the antigen-enriched polyclonal antibody. The proposed sequence is shown below the highlighted regions, and the bottom parts are short peptides corresponding to elements of the sequences. Green lines are peptides generated from the digestion with Chymotrypsin; light orange is LysC, blue is Trypsin, black is Pepsin, and dark orange is Asp-N. **Fig. 2C** illustrates a typical “middle-down” peptide allowing in this particular case, the assembly of a CDR2 with a CDR3 within the same chain. **Fig. 2D** illustrates the resulting peptides from the action of a given protease on a specific germline (IGHV3-53*01) under non-reduced conditions. The black arrows represent possible pepsin sites, the blue ones are peptides resulting from the digestion with trypsin, and the orange ones are peptides from the protease Asp-N. **Fig. 2E** and **2F** illustrate some disulfide-linked peptide analysis spectra. Those two peptides allow pairing each a specific CDR1 sequence with a specific CDR3 sequence.

### Mass Spectrometry and Database Searching

Proteomics data processing involved enriching 3ml plasma with protein G by gravity flow, isolating 22.2 mg of IgG. SARS-CoV-2 spike protein receptor binding domain (RBD) binding experiments were performed using 20mg of this IgG and streptavidin beads coupled with photocleavable biotin. Approximately 190 μg per 3ml plasma was enriched through acid elution, and an additional 10% antibody yield was obtained through UV cleavage.

Parallel protease digestions allow the production of overlapping peptides. Five proteases— Trypsin, LysC, AspN, Chymotrypsin, and Pepsin were used to digest the sample. The sample was split into two sub-samples. The first sample was reduced and alkylated with iodoacetamide. In the second sample, the cysteine residues were converted into a lysine analog using 2-bromoethylamine hydrobromide, rendering the cysteine residues susceptible to cleavage by trypsin and LysC enzymes (23). The digested samples were then analyzed on an Orbitrap Exploris 240 instrument and compared to the NGS IgSeq dataset.

The Mass spectrometry data were searched against the IgSeq dataset. Sequences containing two or more confident peptide-spectrum matches were selected and grouped according to their CDR3 similarities. Sequences of which the CDR3 regions differ by at most one amino acid are grouped together. This results in 183 heavy clusters, 203 kappa clusters, and 290 lambda clusters, respectively (Dataset S1). The majority of these sequences were matched in the framework regions, and only a handful contain full MS coverage of CDR3, including only 4 heavy clusters, 9 kappa clusters, and 4 lambda clusters considered in this study.

### De Novo Sequencing

Following the procedure described in the Materials and Methods section, *de novo* sequencing allowed the extraction of additional sequences not found in the IgSeq chain clusters. This resulted in 3 new heavy chain clusters and 4 new lambda chain clusters with different CDR3s compared to the IgSeq clusters and partial sequences covering different CDR regions. The selected IgSeq sequences and the full and partial *de novo* sequences are available in Dataset S2.

Fig. 2B exemplifies short contigs with distinct peptide overlaps in three CDR regions. The left sequence in Fig. 2B represents a *de novo* CDR1 region with possible neighboring sequences (”CTVSGGSLGRGSDFWGW” in Dataset S2). The center of Fig. 2B presents a CDR2 sequence (”WLGSLYYSGSTTYYNPSLK”), while the right of Fig. 2B displays a CDR3 sequence (”YHCARKQWLRGAFDLWGQG”). Overlapping peptides provide additional confidence in these proposed sequences. Fig. 2C shows a peptide-spectrum match (PSM) between a middle-down MS/MS spectrum and an extended assembly covering significant portions of the CDR2 and CDR3 regions shown above. This spectrum confirms that this pair of CDR2 and CDR3 regions should belong to the same antibody.

The disulfide bond between the two Cysteine residues near CDR1 and CDR3 was used to identify certain pairs of CDR1 and CDR3 (as shown in Fig. 2D). Fig. 2E and Fig. 2F display two examples identified by pLink software using our MS/MS data under non-reducing conditions. *SI Appendix Table S2* lists non-reduced peptides found in this study. Some peptides in *SI Appendix Table S2* confirm the initial entry assembly from Dataset S2. However, extended digestion under neutral conditions may cause disulfide bonds to scramble (24) and generate false positives, though these are usually associated with low-intensity peptides. Therefore, combining the disulfide bond information with other evidence is necessary to determine the CDR pairing definitively.

The separation experiment, performed under non-reduced conditions, provides additional pairing information. We used IdeS digestion and a Native gel to separate antibodies by charge and mass. This method yielded 12 bands (Fig. 3A), subsequently subjected to Trypsin digestion. Fig. 3B shows an example of the normalized quantification vectors of peptides for selected CDR contigs (sequences in *SI Appendix Table S3*). The curves for HCDR1-c02, HCDR2-c01, and HCDR3-c01 show similarity, suggesting they belong to the same antibody protein and should be assembled as part of the same chain. Pearson correlation analysis confirms the highest similarity scores between HCDR1-c02, HCDR2-c01, and HCDR3-c01, supporting their assembly as components of the same chain. Co-elution profile analysis also enables the pairing of a heavy chain with its cognate light chain. Orthogonal separation could improve pairing. If heavy and light chain pairing is uncertain, multiple combinations of heavy and light chains can be generated and tested.

**Figure 3:**
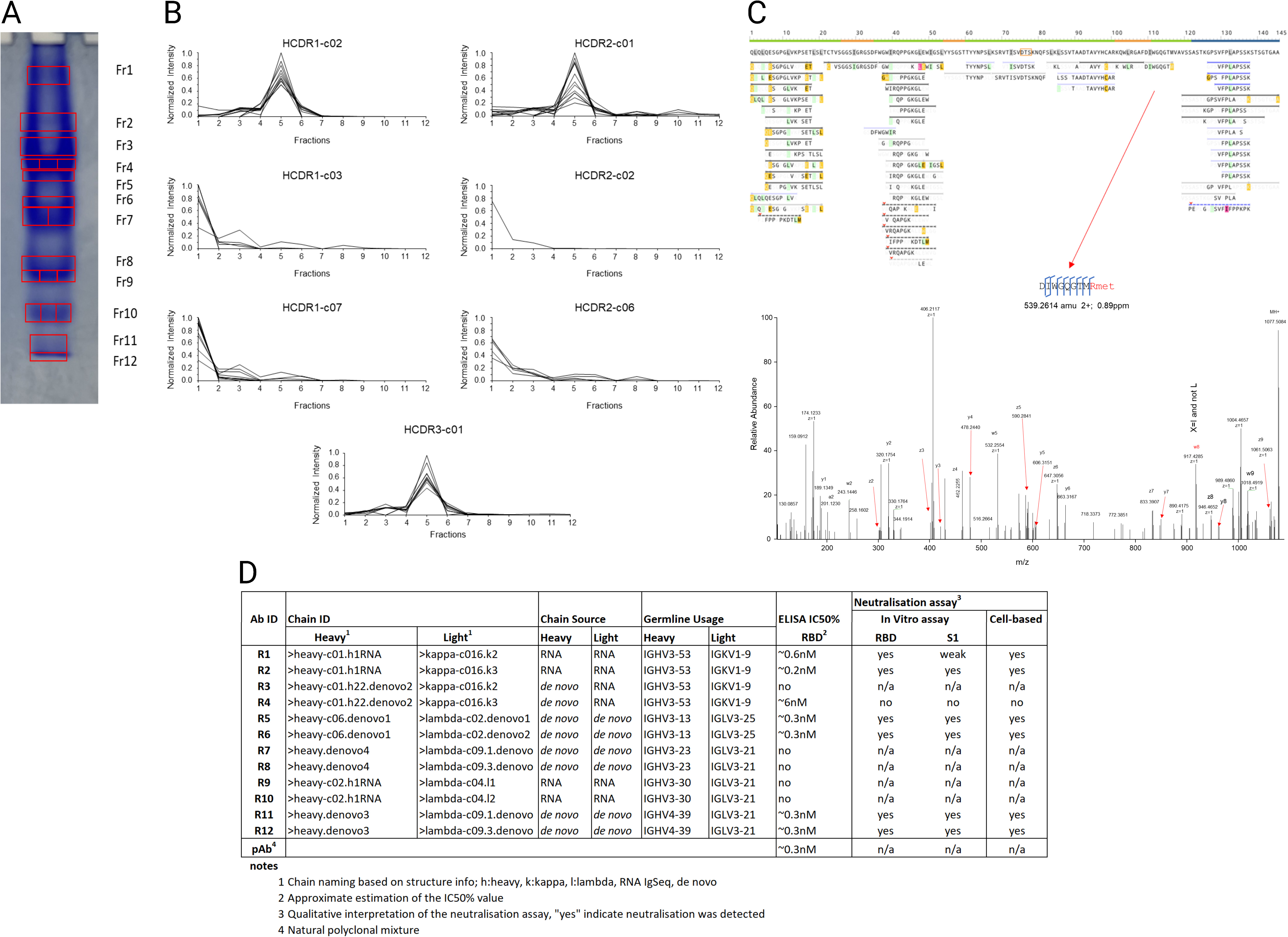
Final Step in Sequencing: CDRs Assembly, Isobaric Ambiguity Resolution, and Recombinant Antibody Testing **Fig. 3A** depicts the separation of the polyclonal antibody on a native gel. Initially, the enriched polyclonal antibody was digested with IdeS, and the resulting sample was fractionated into 12 fractions and further digested with trypsin. **Fig. 3B** illustrates the quantitative profiling of various peptide “contigs” across those 12 fractions, where contigs were grouped based on their similarity to assemble chains and pair heavy-light (HL) chains (more information on the contig sequence can be found in SI Table S3). **Fig. 3C** top: shows an example of the sequence coverage resulting from EThcD analysis of trypsin and pepsin digestion. Peptides resulting from pepsin digestion were labeled with Arginine methyl ester. Bottom: displays EThcD spectra acquired to resolve specific Isoleucine/Leucine assignments; in that case, the generation of w ions under EThcD fragmentation allows the assignment of w8 to an isoleucine. Additionally, in this case, a positive charge (i.e., Methyl Arginine as a y1 ion) was added to the C-terminal end of the non-tryptic peptide to facilitate the C-terminal fragment to be detectable in MSMS mode. **Fig. 3D** shows the expressed candidates and their affinities in ELISA Assay, *in vitro* neutralization, and *in vivo* neutralization assay.

In addition, we used semi-denatured protein on a gel to separate the antibodies and the Fab2 fragments (*SI Appendix* Fig. S1A). Seven heavy chains, nine kappa chains, and six lambda chains were correlated using Native and semi-denaturing gel data. *SI Appendix Fig. S1B* profile correlations showed that five heavy chains might pair with unique light chains. The remaining two heavy chains were associated with weak MS signals and unrelated to the light chains. Additionally, ten light chains had no significant correlation with heavy chains and were excluded from the analysis. Some sequences were ambiguous and required multiple H/L combinations.

Fig. 3C shows the EThcD-based isoleucine (I) and leucine (L) residue assignment method we used in this study, as described by Zhokhov et al. (25). This method produces z- and w-ions from the C-terminus. The w-ions help distinguish Isoleucine and Leucine residues. This method works nicely for Tryptic and LysC digests. Other proteases like Pepsin or Chymotrypsin may produce inconsistent results. To overcome this limitation, we have developed a method that introduces a positive charge on the C-terminal amino acid. This modification generates z-ions to w-ions and helps sequence and identify Isoleucine or Leucine residues. Fig. 3C, bottom part, shows a spectrum of an isoleucine-specific w8 ion detected using this method. Most I/L ambiguities can be resolved using this labeling technique, EThcD, germline analysis, and IgSeq data.

### Sequencing Results and Functional Assays

Dataset S3 contains the final heavy-light pairing results and antibody sequences. The twelve resulting antibodies (R1–R12) are combinations of six heavy and eight light chains. All six heavy chains contain different CDR3. Four of the six chains are absent from IgSeq data and constructed by *de novo* sequencing only. The eight light chains belong to four different clusters. Sequences within each cluster share highly similar CDR3 and V-region sequences. Four light chain sequences (two clusters) are from *de novo* sequencing. Together, six of the 12 antibodies have both heavy and light chains constructed *de novo*, while eight have at least one chain constructed *de novo*.

We expressed the 12 recombinant antibodies and tested their functions with the following assays:

- ELISA and SPR analyses for affinity to RBD domain.
- *In vitro* neutralization assays against RBD and S1 domains, respectively.
- In-vivo neutralization assays against Fluc-GFP pseudovirus.

The results are summarized in Fig. 3D.

ELISA assays were used to assess recombinant antibody affinity for the Receptor-Binding Domain (RBD) (Fig. 4A). Seven of the 12 recombinant antibodies (R1, R2, R4, R5, R6, R11, R12) exhibited high affinities towards the RBD. Six recombinant antibodies have affinities similar or better against the RBD than the natural polyclonal anti-RBD antibody named PD124 (with an IC50 at 0.3-0.6 nM Fig. 4A red arrows). Noticeably, four of the six strongest binders (R5, R6, R11, R12) were obtained by *de novo* sequencing.

**Figure 4:**
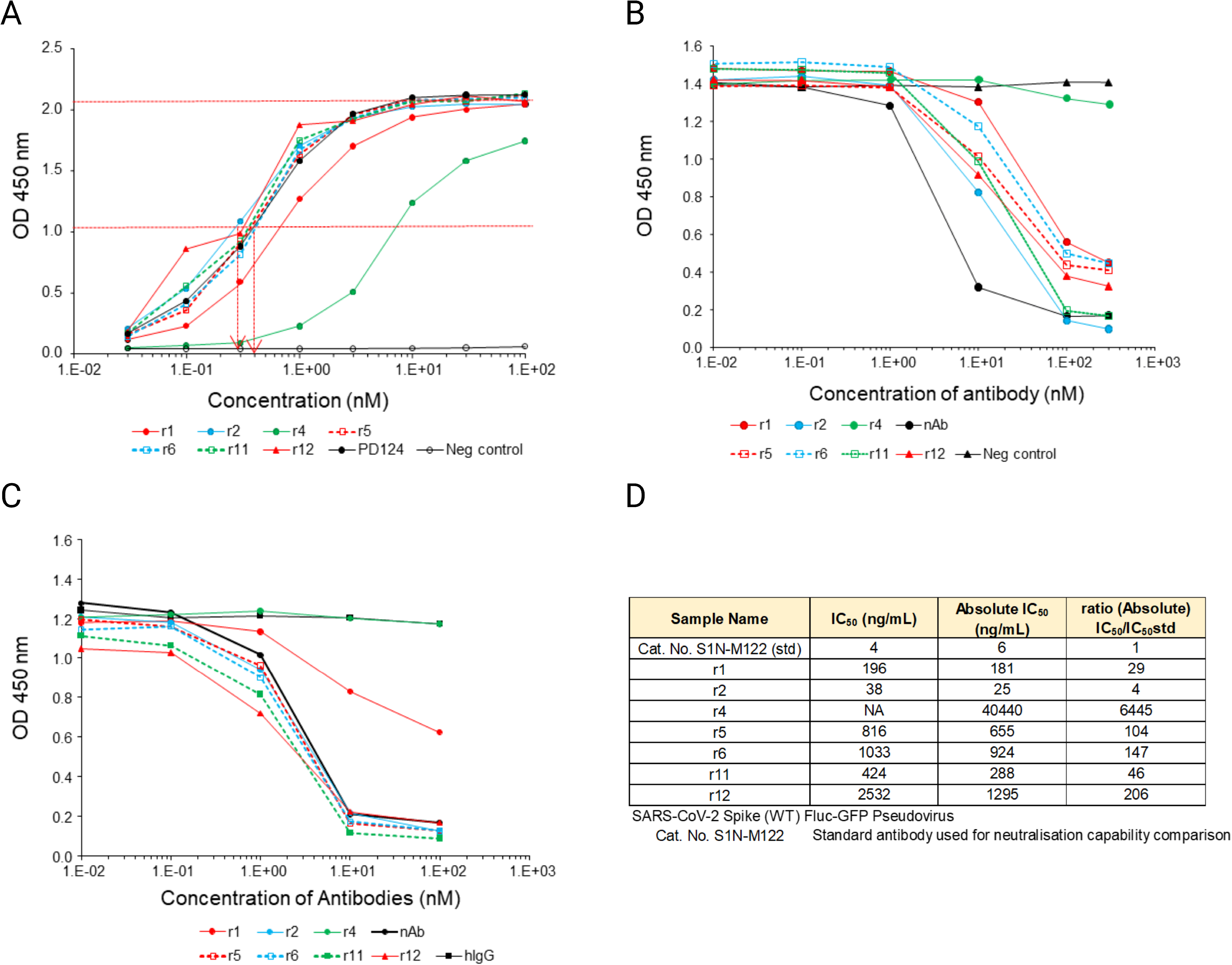
Performance and properties of the investigated antibodies. The Supporting Materials and Methods section describes how the various ELISA procedures were performed. **Fig. 4A** presents a conventional ELISA assay comparing recombinant antibodies with the natural polyclonal antibody PD124 enriched in this study as a benchmark measure. The red dashed arrows are the IC50 for the best recombinant antibody (R2 at ca 3nM) versus the natural polyclonal antibody (PD124 at approx. 4nM). Nonbinding recombinant antibodies (r3, r7, r8, r9, r10) were eliminated from further comparisons. Due to its restricted availability, the natural polyclonal antibody PD124 was excluded from other investigations. **Fig. 4B** illustrates a pseudo-neutralization assay conducted on the recombinant forms found in **Fig. 4A**, utilizing the RBD domain. **Fig. 4C** is also a pseudo-neutralization assay that targets the S1 protein as the bait in the test. **Fig. 4D** displays an in vivo neutralization test that utilizes SARS-CoV-2-Spike/Flu-GFP pseudo-virus to neutralize a positive standard S1N-M122. The assay measures the ratios of Absolute IC50.

The seven binders were further assessed using Surface Plasmon Resonance (SPR) methods, including complex stability analysis (26) and off-rate screening (27). Complex stability, which indicates the fidelity between the antibody *and antigen, was evaluated by plotting the stability early vs. stability late values (SI Appendix Fig. S2A-S2C*). This analysis showed that R11 and R12 bind with the highest complex stability, followed by R2, R6, and R5. The off-rate screening was performed as a complementary analysis to further investigate the top candidates’ kinetic properties. Multiparametric evaluation was completed with this method. The top binders (R2, R5, R6, R11, and R12) were categorized with slow off-rates (*SI Appendix Fig. S2*D), indicating their prolonged binding capabilities. The dissociation rates of R4 and R1 were much faster in comparison.

*In vitro*, pseudo-neutralization assays were performed on the seven binders to determine if they could interfere with the ACE/RBD and ACE/S1 interactions (Fig. 4B and Fig. 4C, respectively). Due to its limited availability, the natural affinity-purified polyclonal antibody (PD124) was not used as a neutralization benchmark. Among the tested antibodies, R1, R2, R5, R6, R11, and R12 exhibited neutralization capability, while R1 demonstrated reduced neutralization ability against S1 compared to the RBD protein. R4 did not show any significant neutralization capability.

A cell-based neutralization assay was performed to test those seven recombinant antibodies’ neutralization activities against Fluc-GFP pseudovirus (Fig. 4D). All tested recombinant antibodies displayed varying degrees of neutralization activities. Relative to the positive control S1N-M122, R2 showed the closest neutralization efficacy, followed by R1 and R11.

## Discussion

The majority (approximately 90%) of human antibody discovery related to SARS-CoV was achieved through B-cell sequencing-only approaches, as evidenced by a survey of the CoV-AbDab database (SI Appendix Fig. S1C). Although this approach has been successful, there is often limited correspondence between the identified B-cells and the circulating IgG population, indicating the potential for missed valuable candidates (16, 18). It should be noted that IgG molecules in circulation, rather than the B-cells themselves, serve as the final product and primary effectors responsible for humoral immunity. Most of the circulating IgGs are produced by large plasma cells in the bone marrow rather than circulating B-cell (28). While combined proteomics and B-cell repertoire analysis can confirm the presence of specific antibodies in circulation, it cannot identify antibodies that may have been missed by B-cell sequencing. *De novo* pAb sequencing, although technically challenging, offers the most promising approach to comprehensively understanding the final products of the humoral immune response.

This present study employed a combination of conventional proteomics and *de novo* sequencing approaches. Our *de novo* sequencing efforts led to the discovery of six new antibodies, where both the heavy and light chains were absent from IgSeq data. Four of these six antibodies demonstrated equivalent affinity to the natural pAb, while only two IgSeq antibodies showed similar affinity levels. Additionally, the *de novo* and IgSeq antibodies exhibited varying degrees of neutralization activity for both *in vitro* and cell-based neutralization assays. The neutralizing activities against the Fluc-GFP pseudovirus were comparable to those previously reported by He et al. (29). It is important to note that neutralization depends on binding to the correct epitope, resulting in variations in neutralization abilities even among mAbs with similar binding affinities (30). The demonstrated binding affinity and neutralizing activities strongly support the accuracy of our sequences. Furthermore, our findings indicate that the circulating B-cell repertoire does not encompass all sequences of circulating antibodies, underscoring the necessity of *de novo* sequencing.

In the study by Guthals et al. (17), 28 antibodies were sequenced and expressed, but only two exhibited binding potencies, albeit significantly lower than the original pAb, indicating potential sequence errors. Notably, due to the absence of a pairing method for heavy and light chains, all combinations of four heavy and seven light chains had to be expressed and tested. More recently, Bondt et al.(22, 31) attempted *de novo* sequencing of a polyclonal antibody using a similar approach as in Guthals et al. (17). However, no binding or neutralization data were provided to validate the accuracy or effectiveness of the generated antibodies. In our experience, numerous possible sequences exist for each CDR, resulting in millions of potential combinations for an antibody’s six CDRs. Resolving the assembly ambiguity necessitates more comprehensive information than overlapping peptides, middle-down MS datasets, and intact mass data acquisition. Consequently, we suspect CDRs from different mAbs may have been incorrectly assembled in the works mentioned above, leading to suboptimal binding outcomes. A significant distinction in our approach lies in extensively utilizing various experiments (e.g., middle-down MS, non-reducing digestion, and different antibody separation techniques) to generate orthogonal datasets. We combined these orthogonal datasets to assemble the different CDRs into functional antibodies accurately.

In addition to previous studies (7, 8, 17, 22, 32), we have developed a method for pairing heavy and light chains to produce functional antibodies. Using bulk sequencing data from B cells is challenging as the heavy and light chain sequences are not paired. Previous studies attempted all possible pairings (7, 8, 17), but this becomes impractical and expensive when there are more than 10 heavy and light chains. Single B-cell sequencing provides pairing information but is more laborious and costly than bulk sequencing. Additionally, the limited number of cells in a single-cell sequencing experiment reduces the coverage of the B cell repertoire. Utilizing a co-separation MS profile for pairing, our approach offers an alternative solution to overcome these challenges.

Furthermore, we implemented stringent validation measures to confirm the functionality of the assembled antibodies. Extensive binding assays and neutralization tests assessed their functional activities. These thorough experimental validations provide solid evidence for the correctness and effectiveness of our approach.

*De novo* sequencing is vital when B cells cannot be obtained. For instance, immunized animals produce most of the current pAb reagents. However, suppose the production of the same effective polyclonal antibody ceases in subsequent immunization rounds or is no longer available. In that case, de novo sequencing of stored protein samples can still recover the original polyclonal antibody. Our unpublished data provides evidence for the efficacy of this approach in multiple instances.

While previous studies have demonstrated successful *de novo* sequencing of a single protein or monoclonal antibody (mAb) (19, 20, 32), it is essential to note that *de novo* sequencing of a polyclonal antibody (pAb) poses more significant challenges. Furthermore, antibodies within a pAb mixture often share similar framework regions, making it challenging to pair CDRs using peptide overlaps unambiguously. The high sequence diversity within CDRs results in diluted peptide signals, reducing the signal-to-noise ratio and further complicating *de novo* sequencing. Coincidentally, CDRs are the most critical regions of antibodies. Additionally, not all peptides produce equally strong mass spectrometry (MS) signals (33, 34), potentially leading to a failure in sequencing antibodies if even one CDR is missing from the MS data. In our study, we successfully *de novo* sequenced more CDR sequences than the final list of antibodies, highlighting the challenges involved (Dataset S2). Additional fractionation of the pAb could improve MS coverage. Still, it would also increase the dataset size and the complexity of handling the more significant number of CDR combinations, requiring more advanced algorithms.

Although the number of reported antibodies in our study may appear small, numerous partial sequences in Dataset S2 suggest that the initial polyclonal antibody (pAb) sample is a highly complex mixture, even after antigen affinity enrichment. This circumstance highlights the difficulty of our pAb *de novo* sequencing approach and the potential for sequencing a larger number of antibodies with further methodological improvements. It is worth noting that intense mass spectrometry (MS) signals are typically generated from peptides resulting from the protease digestion of more abundant proteins, suggesting that the most prevalent antibodies in the pAb mixture are likely to be among the first successfully *de novo* sequenced.

Despite the challenges mentioned above, our study has successfully demonstrated the feasibility of *de novo* sequencing from a human serum polyclonal antibody (pAb), generating multiple monoclonal antibodies (mAbs) that closely resemble the original pAb. This achievement emphasizes the potential of pAb *de novo* sequencing as a valuable strategy for harnessing the natural immune response of animals and humans, enabling the discovery of novel antibody reagents and therapeutics. We anticipate that with further advancements in mass spectrometry experiments and bioinformatics, the field of pAb *de novo* sequencing will continue to flourish and contribute to antibody research and development.

## Materials and Methods

A comprehensive Materials and Methods section can be found in the SI Appendix.

### Sample collection

Discovery Life Sciences collected three healthy donor samples; In this study, we present data specifically from one donor, labeled as “522”. The individual is a 49-year-old Caucasian female, and further details can be found in SI Appendix Table S1. PBMC were collected from blood using BD Vacutainer CPT™ following supplier instructions and stored in Qiagen RNAprotect® Cell Reagent. We collected proteomics samples: After centrifuging blood, 50% was transferred to a Matrix cryo vial.

### RNA Extraction

PBMCs were lysed and homogenized using a QIA shredder (Qiagen) before RNA extraction following the supplier’s recommendation. PBMC and RNAprotect Cell Reagent were centrifuged to isolate the cell pellet. Beta-mercaptoethanol was added to the cell pellet and mixed. The lysate was centrifuged in a QIA shredder spin column. RNA extraction was performed on the homogenized lysate from the QIA shredder flow-through in a new tube. PBMC total RNA was isolated using the RNeasy Mini Kit (Qiagen) per the manufacturer’s instructions.

### cDNA synthesis, BCR Amplification, and Sequencing Library Generation

The SMARTer Human BCR IgG IgM H/K/L Profiling Kit (TakaraBio) was used to construct libraries for NGS. To generate cDNA, one µg of RNA was input into the first strand cDNA synthesis reaction following instructions from the BCR profiling kit. PCR conditions for BCR amplification and sequencing library generation were followed according to the kit user manual.

### Next-Generation Sequencing

The final products were re-quantified using the 1X dsDNA HS assay (Thermo Fisher Scientific). The final quantified libraries were normalized and pooled to a 4nM library, which was used to prepare an 8 pM loading library. The Illumina MiSeq reagent kit v3 was used to sequence 600 cycle paired-end reads.

### IgG enriched against the antigen

Total IgG was from 3 mL human serum from a patient vaccinated against SARS-CoV-2 using protein G agarose resin (Genscript). Protein G resin was washed with PBS and eluted with 0.1M glycine buffer, pH 2.5. After eluting IgG, the buffer was exchanged into PBS using an Amicon filter (Sigma-Aldrich). Affinity enriched: SARS-CoV-2 spike protein Receptor Binding Domain, RBD (ExonBio) was coupled to streptavidin-coated agarose beads (Sigma-Aldrich) to enrich anti-RBD antibodies. Then, 20 mg of total IgG was added, incubated, and washed. Then, the anti-RBD antibodies were eluted with glycine pH 2.5.

### The in-solution protease digestion procedure

was performed with two different cysteine modifications. Using iodoacetamide with Trypsin, LysC, AspN, pepsin, chymotrypsin, and with 2-bromoethylamine hydrobromide, BEA, converting the cysteine into a lysine-like amino acid(23). This last sample was digested with trypsin and LysC.

### Protein Separation

Native gel: The affinity-enriched fraction, PD124, underwent separation on Biorad precast 7.5% polyacrylamide gels with and without IdeS. Excised bands were washed and divided into 12 segments (Fig. 3A). Each band was subjected to trypsin digestion using a standard in-gel protocol (35). NRT (Non-reducing Room Temperature) gel (*SI Appendix* Fig. S1A): PD124 underwent separation using a modified non-reducing SDS-PAGE for heavy-light chain pairing without pre-heating. Affinity-enriched IgG, either in its native form or after IdeS digestion, was loaded at 10 µg per lane on Biorad precast 7.5% Mini-PROTEAN TGX gels. Each band was treated with trypsin using a standard in-gel protocol (35).

### Digestion under the non-reducing conditions

The affinity-enriched polyclonal antibody was dried completely under low pressure (i.e. Speedvac), then reconstituted in 8 M urea in 100 mM tris buffer, then N-ethylmaleimide (NEM) was added to block all free cysteine. The endoprotease LysC was initially used, followed by digestion with AspN. After protease digestion, the sample was dried under low pressure and reconstituted in 40 µL of 0.1% formic acid. Next, 2.5 µL of digested samples were loaded onto Evotips per the manufacturer’s instructions.

### Isobaric resolution

Three selected samples were analyzed in EThCD mode to resolve potential isobaric ambiguity (such as isobaric amino acid I/L). These samples are the previous trypsin and pepsin digest and were used as follow: the trypsin was used as is and the pepsin one was labeled at the C-terminal end using Arginine methyl ester dihydrochloride (Sigma-Aldrich) using the method used in (36).

### Mass spectrometry

HCD mode: Samples from in-solution, in-gel, and non-reduced digestion were all run on Orbitrap 240 Exploris (Thermo Fisher Scientific) using 30 samples per day, 44 min method on a 15cm PepSep column. EThcD mode: An Orbitrap Eclipse Tribid Mass spectrometer (Thermo Fisher Scientific) connected to an Evosep operated in a 30 samples per day was used to resolve the isobaric ambiguities.

### Database Searching

The NGS reads were translated into protein sequences to create an antibody sequence database, then matched with mass spectrometry data using Novor.Cloud (http://novor.cloud). The antibody sequences were sorted based on MS coverage in CDR regions and total sequence coverage.

### De Novo Sequencing

The MS/MS spectra were first *de novo* sequenced using Novor software to obtain peptide sequences (37). If two *de novo* peptide sequences overlap and the overlapping region contains at least two mutated amino acids compared to the germline, the two sequences are considered from the same antibody and merged to form a longer sequence contig. This continues until all pairs are gone. Aligning contigs with germline genes yields candidate sequences for each CDR. Longer contigs may span multiple CDRs. These longer contigs spanning multiple CDRs were filtered by two criteria to ensure quality: they either have a confident middle-down MS/MS spectrum spanning two CDRs, or their peptide overlaps are prominent (containing three or more mutations for each overlap). SI Appendix Table S3 displays assembly contig sequences. The peak area of each unique peptide in CDR contigs from each fraction in NRT and native gel separation experiments was calculated using MaxQuant software (Ver2.1.3). To quantify peptides, a normalized quantification vector was generated for each separation experiment by dividing the quantity of each peptide in each fraction by the total quantity in all fractions. The similarity score between contigs was determined as the average Pearson correlation coefficient between every pair of unique peptides from the two contigs. The similarity scores from multiple separation experiments were combined. CDR contigs with high total similarity scores, covering CDR1, CDR2, and CDR3, were assembled together, indicative of being from the same antibody. The software pLink (38) was employed to identify disulfide bridge-containing peptides through non-reduced protease digestion, aiding in eliminating quantification ambiguities in contig assembly. The assembled contigs were aligned with the germline sequence, and any potential gaps in the framework region were addressed by augmenting with overlapping de novo peptides, thus achieving a full chain sequence.

### Heavy-Light Pairing

The lists of heavy and light chains obtained from database searching and *de novo* sequencing were combined for pairing. The similarity score between heavy and light chains was calculated by averaging the Pearson correlation scores of unique peptides. High-scoring pairs were considered candidate antibodies. MS coverage and similarity scores were evaluated empirically to rank the candidate clones. The sequences can be found in Dataset S3.

The details regarding the different ELISA and SPR assays performed in this study are available in the Materials and Methods in the SI Appendix.

## Notes

### Competing Interest Statement

The authors disclose the following competing financial interests: BM and QL have equity interests in Rapid Novor, a company that may potentially benefit from the research outcomes. All other authors are employees of Rapid Novor.

## References

1. R. M. Lu, et al., Development of therapeutic antibodies for the treatment of diseases. J Biomed Sci 27 (2020).

2. B. Hjelm, B. Forsström, J. Löfblom, J. Rockberg, M. Uhlén, Parallel Immunizations of Rabbits Using the Same Antigen Yield Antibodies with Similar, but Not Identical, Epitopes. PLoS One 7 (2012).

3. G. Kohler, C. Milstein, Continuous cultures of fused cells secreting antibody of predefined specificity. Nature 256, 495–497 (1975).

4. P. C. Wilson, S. F. Andrews, Tools to therapeutically harness the human antibody response. Nat Rev Immunol 12, 709–719 (2012).

5. A. Frenzel, T. Schirrmann, M. Hust, Phage display-derived human antibodies in clinical development and therapy. MAbs 8, 1177–1194 (2016).

6. A. Pedrioli, A. Oxenius, Single B cell technologies for monoclonal antibody discovery. Trends Immunol 42, 1143–1158 (2021).

7. W. C. Cheung, et al., A proteomics approach for the identification and cloning of monoclonal antibodies from serum. Nat Biotechnol 30, 447–452 (2012).

8. S. Sato, et al., Proteomics-directed cloning of circulating antiviral human monoclonal antibodies. Nat Biotechnol 30, 1039–1043 (2012).

9. J. Lavinder, Y. Wine, D. Boutz, E. Marcotte, G. Georgiou, PROTEOMIC IDENTIFICATION OF ANTIBODES. 1–87 (2012).

10. G. Georgiou, et al., The promise and challenge of high-throughput sequencing of the antibody repertoire. Nat Biotechnol 32, 158–168 (2014).

11. R. K. A. França, et al., Progress on Phage Display Technology: Tailoring Antibodies for Cancer Immunotherapy. Viruses 15 (2023).

12. E. Doevendans, H. Schellekens, Immunogenicity of innovative and biosimilar monoclonal antibodies. Antibodies 8 (2019).

13. S. J. Zost, et al., Potently neutralizing and protective human antibodies against SARS-CoV-2. Nature 584, 443–449 (2020).

14. K. Westendorf, et al., LY-CoV1404 (bebtelovimab) potently neutralizes SARS-CoV-2 variants. Cell Rep 39 (2022).

15. I. S. Georgiev, et al., Delineating antibody recognition in polyclonal sera from patterns of HIV-1 isolate neutralization. Science (1979) 340, 751–756 (2013).

16. N. Chaudhary, D. R. Wesemann, Analyzing immunoglobulin repertoires. Front Immunol 9 (2018).

17. A. Guthals, et al., De Novo MS/MS Sequencing of Native Human Antibodies. J Proteome Res 16, 45–54 (2017).

18. J. Chen, et al., Proteomic Analysis of Pemphigus Autoantibodies Indicates a Larger, More Diverse, and More Dynamic Repertoire than Determined by B Cell Genetics. Cell Rep 18, 237–247 (2017).

19. N. Bandeira, V. Pham, P. Pevzner, D. Arnott, J. R. Lill, Automated de novo protein sequencing of monoclonal antibodies. Nat Biotechnol 26, 1336–1338 (2008).

20. X. Liu, et al., De Novo protein sequencing by combining top-down and bottom-up tandem mass spectra. J Proteome Res 13, 3241–3248 (2014).

21. W. Peng, M. F. Pronker, J. Snijder, Mass Spectrometry-BasedDe NovoSequencing of Monoclonal Antibodies Using Multiple Proteases and a Dual Fragmentation Scheme. J Proteome Res 20, 3559–3566 (2021).

22. A. Bondt, et al., Into the dark serum proteome: personalized features of IgG1 and IgA1 repertoires in severe COVID-19 patients. Molecular & Cellular Proteomics, 100690 (2023).

23. M. Thevis, R. R. Ogorzalek Loo, J. A. Loo, In-gel derivatization of proteins for cysteine-specific cleavages and their analysis by mass spectrometry. J Proteome Res 2, 163–172 (2003).

24. W. C. Sung, et al., Evaluation of disulfide scrambling during the enzymatic digestion of bevacizumab at various pH values using mass spectrometry. Biochim Biophys Acta Proteins Proteom 1864, 1188–1194 (2016).

25. S. S. Zhokhov, S. V. Kovalyov, T. Y. Samgina, A. T. Lebedev, An EThcD-Based Method for Discrimination of Leucine and Isoleucine Residues in Tryptic Peptides. J Am Soc Mass Spectrom 28, 1600–1611 (2017).

26. P. Säfsten, S. L. Klakamp, A. W. Drake, R. Karlsson, D. G. Myszka, Screening antibody-antigen interactions in parallel using Biacore A100. Anal Biochem 353, 181–190 (2006).

27. J. B. Murray, S. D. Roughley, N. Matassova, P. A. Brough, Off-rate screening (ORS) by surface plasmon resonance. An efficient method to kinetically sample hit to lead chemical space from unpurified reaction products. J Med Chem 57, 2845–2850 (2014).

28. R. L. Longmire, et al., In vitro splenic igg synthesis in hodgkin’s disease. New England Journal of Medicine 289, 763–767 (1973).

29. B. He, et al., Rapid isolation and immune profiling of SARS-CoV-2 specific memory B cell in convalescent COVID-19 patients via LIBRA-seq. Signal Transduct Target Ther 6 (2021).

30. R. Rouet, et al., Broadly neutralizing SARS-CoV-2 antibodies through epitope-based selection from convalescent patients. Nat Commun 14 (2023).

31. A. Bondt, et al., Human plasma IgG1 repertoires are simple, unique, and dynamic. Cell Syst 12, 1131–1143.e5 (2021).

32. X. Liu, Y. Han, D. Yuen, B. Ma, Automated protein (re)sequencing with MS/MS and a homologous database yields almost full coverage and accuracy. Bioinformatics 25, 2174– 2180 (2009).

33. T. Le Bihan, M. D. Robinson, I. I. Stewart, D. Figeys, Definition and characterization of a “trypsinosome” from specific peptide characteristics by Nano-HPLC-MS/MS and in silico analysis of complex protein mixtures. J Proteome Res 3, 1138–1148 (2004).

34. Y. F. Li, R. J. Arnold, H. Tang, P. Radivojac, The importance of peptide detectability for protein identification, quantification, and experiment design in MS/MS proteomics. J Proteome Res 9, 6288–6297 (2010).

35. M. Mann, R. C. Hendrickson, A. Pandey, “ANALYSIS OF PROTEINS AND PROTEOMES BY MASS SPECTROMETRY” (2001).

36. T. Le BIHAN, B. MA, Z. Mcdonald, Q. Liu, P. Taylor, A LABELLING METHOD TO DISTINGUISH ISOBARIC AMINO ACIDS AND AMINO ACID COMBINATIONS. WO/2020/124252.

37. B. Ma, Novor: Real-Time Peptide de Novo Sequencing Software. J Am Soc Mass Spectrom 26, 1885–1894 (2015).

38. S. Lu, et al., Mapping native disulfide bonds at a proteome scale. Nat Methods 12, 329– 331 (2015).

