## Supplementary material for "*de Novo* Sequencing of Antibodies for Identification of Neutralizing Antibodies in Human Plasma Post SARS-CoV-2 Vaccination": Dataset S3

#### Antibodies Sequenced and Expressed:

| Ab | Heavy | Light | De novo |
| --- | --- | --- | --- |
| R1 | >heavy-c01.ARDLQDYGMDV.h1RNA | >kappa-c016.k2.QQLNSYPRT | - |
| R2 | >heavy-c01.ARDLQDYGMDV.h1RNA | >kappa-c016.k3.QQLNSYPRGYT | - |
| R3 | >heavy-c01.h22.ARELGDGMDV.denovo2 | >kappa-c016.k2.QQLNSYPRT | H |
| R4 | >heavy-c01.h22.ARELGDGMDV.denovo2 | >kappa-c016.k3.QQLNSYPRGYT | H |
| R5 | >heavy-c06.ARVGTTGYDLYGMDV.denovo1 | >lambda-c02.QSGDSGGWV.denovo1 | H/L |
| R6 | >heavy-c06.ARVGTTGYDLYGMDV.denovo1 | >lambda-c02.QSGDSGGWV.denovo2 | H/L |
| R7 | >heavy-AKTQWEHEGSDFDY.denovo4 | >lambda-c09.1.QVWDSGTDHSHVV.denovo | H/L |
| R8 | >heavy-AKTQWEHEGSDFDY.denovo4 | >lambda-c09.3.QVWDSGTDHSHVV.denovo | H/L |
| R9 | >heavy-c02.ARYNSGYYALDY.h1RNA | >lambda-c04.l1.QVWDRSSDHPVV | - |
| R10 | >heavy-c02.ARYNSGYYALDY.h1RNA | >lambda-c04.l2.QVWDSSSDVV | - |
| R11 | >heavy.ARKQWLRGAFDI.denovo3 | >lambda-c09.1.QVWDSGTDHSHVV.denovo | H/L |
| R12 | >heavy.ARKQWLRGAFDI.denovo3 | >lambda-c09.3.QVWDSGTDHSHVV.denovo | H/L |

#### Chain Sequences

>heavy-c01.ARDLQDYGMDV.h1RNA_IGHV3-53*05

EVQLVETGGGLIQPGGSLRLSCAASGVTVSSNYMSWVRQAPGKGLEWVSLIYSGGSTFYADSVKGRFTISRDNSKNTLYLQMNSLRVEDTAVYYCARDLQDYGMDVWGQGTTVTVSS**ASTKGPSVFPLAPSSKSTSGTGAAL**

>heavy-c01.h22.ARELGDGMDV.denovo2_IGHV3-53*05

EVQLVETGGGLIQPGGSLRLSCAASGFIVSRNYMSWVRQAPGKGLEWVSVIYSGGSTFYADSVKGRFTISRDNSKNTLYLQMNSLRDEDTAVYYCARELGDGMDVWGQGTTVTVSS**ASTKGPSVFPLAPSSKSTSGGTAAL**

>heavy-c06.ARVGTTGYDLYGMDV.denovo1_IGHV3-13*04

EVQLVESGGDLVQPGGSLRLSCAASGFTFSNYDMHWVRQVTGKGLEWVSGIGKDGDTYYLGSVKGRFAISRDNAKNSLYLQMNSLRAGDTALYYCARVGTTGYDLYGMDVWGQGTTVTVSS**TSTKGPSVFPLAPSSKSTSGGTAAL**

>heavy-AKTQWEHEGSDFDY.denovo4_IGHV3-23*03

EVQLMESGGGLVQPGRSLRLSCAASGFTFDDYAMHWVRQAPGKGLEWVSVIYSGGSSNHADSVKGRFTISRDNSKNTVYLQMNSLRAEDTAVYYCAKTQWEHEGSDFDYWGQGTLVTVSS**ASTKGPSVFPLAPSSKSTSGGTAAL**

>heavy-c02.ARYNSGYYALDY.h1RNA_IGHV3-33*08

QVQLVESGGGVVQPGRSLRLSCAASGFTFSGSGMHWVRHIPGKGLQWVAVIWSDGSNIYYGDSVKGRFTISRDNSKNTLYLQMNSLRAEDTAVYYCARYNSGYYALDYWGRGTLVTVSS**ASTKGPSVFPLAPSSKSTSGTGAAL**

>heavy.ARKQWLRGAFDI.denovo3_IGHV4-39*09

QLQLQESGPGLVKPSETLSLTCTVSGGSIGRGSDFWGWIRQPPGKGLEWIGSLYYSGSTTYYNPSLKSRVTISVDTSKNQFSLKLSSVTAADTAVYHCARKQWLRGAFDIWGQGTMVAVSS**ASTKGPSVFPLAPSSKSTSGTGAAL**

>kappa-c016.k2.QQLNSYPRT_IGKV1-9*01

DIQLTQSPSFLSASVGDRVTITCRASQGISSYLAWYQQKPGKAPKLLIYAASTLQSGVPSRFSGSGSGTEFTLTISSLQPEDFATYYCQQLNSYPRTFGQGTKVEIK**RTVAAPSVFIFPPS**

>kappa-c016.k3.QQLNSYPRGYT_IGKV1-9*01

DIQLTQSPSFLSASVGDRVTITCRASQGISSYLAWYQQKPGKAPKLLIYAASTLQSGVPSRFSGSGSGTEFTLTISSLQPEDFATYYCQQLNSYPRGYTFGQGTKLEIK**RTVAAPSVFIFPPS**

>lambda-c02.QSGDSGGWV.denovo1_IGLV3-25*03

SYELTQPPSVSVSPGQTARITCSGNVFPRQYAYWYQQKPGQAPVLLIYKDSERPSGIPERFSGSGSGTTVTLTITGVQAEDEADYYCQSGDSGGWVFGGGTKLTVL**GQPKAAPSVTLFPPSS**

>lambda-c02.QSGDSGGWV.denovo2_IGLV3-25*03

SYELTQPPSVSVSPGQTARITCSGDALPKQYAYWYQQKPGQAPVLLIYKDSERPSGIPERFSGSGSGTTVTLTITGVQAEDEADYYCQSGDSGGWVFGGGTKLTVL**GQPKAAPSVTLFPPSS**

>lambda-c09.1.QVWDSGTDHSHVV.denovo_IGLV3-21*02

VYLTQPPSVSVAPGQTARITCGGNNIGSKNVHWYQQKPGQAPVLVVFEDSDRPSGIPDRFSGSNSGNTATLTISRVEAGDEADYYCQVWDSGTDHSHVVFGGGTKLTVL**GQPKAAPSVTLFPPSS**

>lambda-c09.3.QVWDSGTDHSHVV.denovo_IGLV3-21*02

VYLTQPPSVSVAPGQTARISCGGNEIGSKSVHWYQQKPGQAPVLVVFDDSDRPSGIPERFSGSNSGNTATLTISRVEAGDEADYYCQVWDSGTDHSHVVFGGGTKLTVL**GQPKAAPSVTLFPPSS**

>lambda-c04.l1.QVWDRSSDHPVV_IGLV3-21*02

SYVLTQPPSVSVAPGQTARITCGGNNIGSRNLHWYQQKPGQAPVLVVNDDSDRPSGIPERFSGSNSGNTATLTINRVEAGDEADYYCQVWDRSSDHPVVFGGGTKLTVL**GQPKAAPSVTLFPPSS**

>lambda-c04.l2.QVWDSSSDVV_IGLV3-21*02

SYVLTQPPSVSVAPGQTARITCGGNNIGSKSVHWYQQKPGQAPVLVVYDDSDRPSGIPERFSGSNSGNTATLTISRVEAGDEADYYCQVWDSSSDVVFGGGTKLTVL**GQPKAAPSVTLFPPSS**

>Heavy_constant

ASTKGPSVFPLAPSSKSTSGGTAALGCLVKDYFPEPVTVSWNSGALTSGVHTFPAVLQSSGLYSLSSVVTVPSSSLGTQTYICNVNHKPSNTKVDKKVEPKSCDKTHTCPPCPAPELLGGPSVFLFPPKPKDTLMISRTPEVTCVVVDVSHEDPEVKFNWYVDGVEVHNAKTKPREEQYNSTYRVVSVLTVLHQDWLNGKEYKCKVSNKALPAPIEKTISKAKGQPREPQVYTLPPSRDELTKNQVSLTCLVKGFYPSDIAVEWESNGQPENNYKTTPPVLDSDGSFFLYSKLTVDKSRWQQGNVFSCSVMHEALHNHYTQKSLSLSPGK

>Kappa_constant

RTVAAPSVFIFPPSDEQLKSGTASVVCLLNNFYPREAKVQWKVDNALQSGNSQESVTEQDSKDSTYSLSSTLTLSKADYEKHKVYACEVTHQGLSSPVTKSFNRGEC

>Lambda_constant

GQPKAAPSVTLFPPSSEELQANKATLVCLISDFYPGAVTVAWKADSSPVKAGVETTTPSKQSNNKYAASSYLSLTPEQWKSHRSYSCQVTHEGSTVEKTVAPTECS

### Coverage Map

**>heavy-c01.ARDLQDYGMDV.h1RNA_IGHV3-53*05**

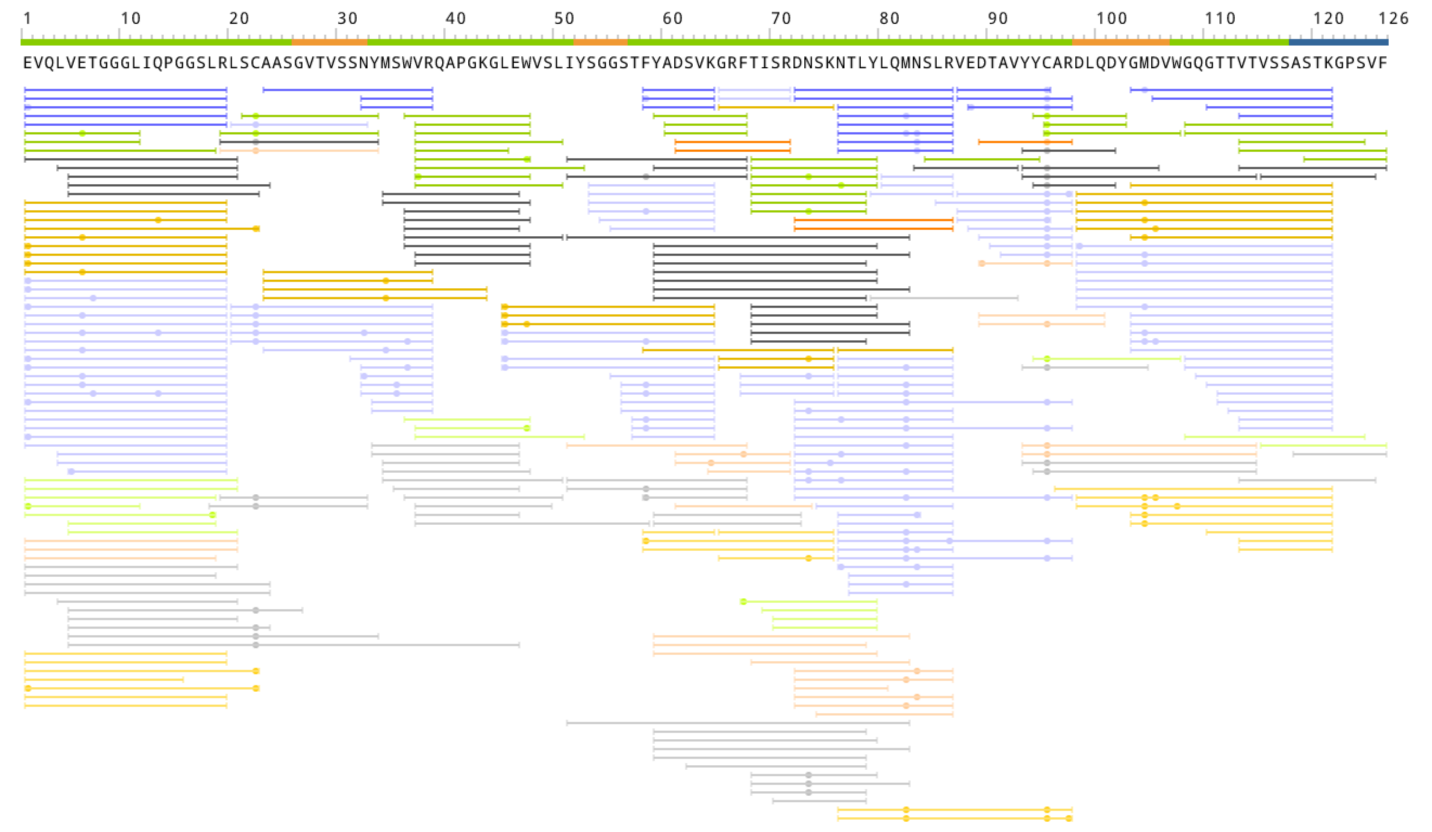

**>heavy-c01.h22.ARELGDGMDV.denovo2_IGHV3-53*05**

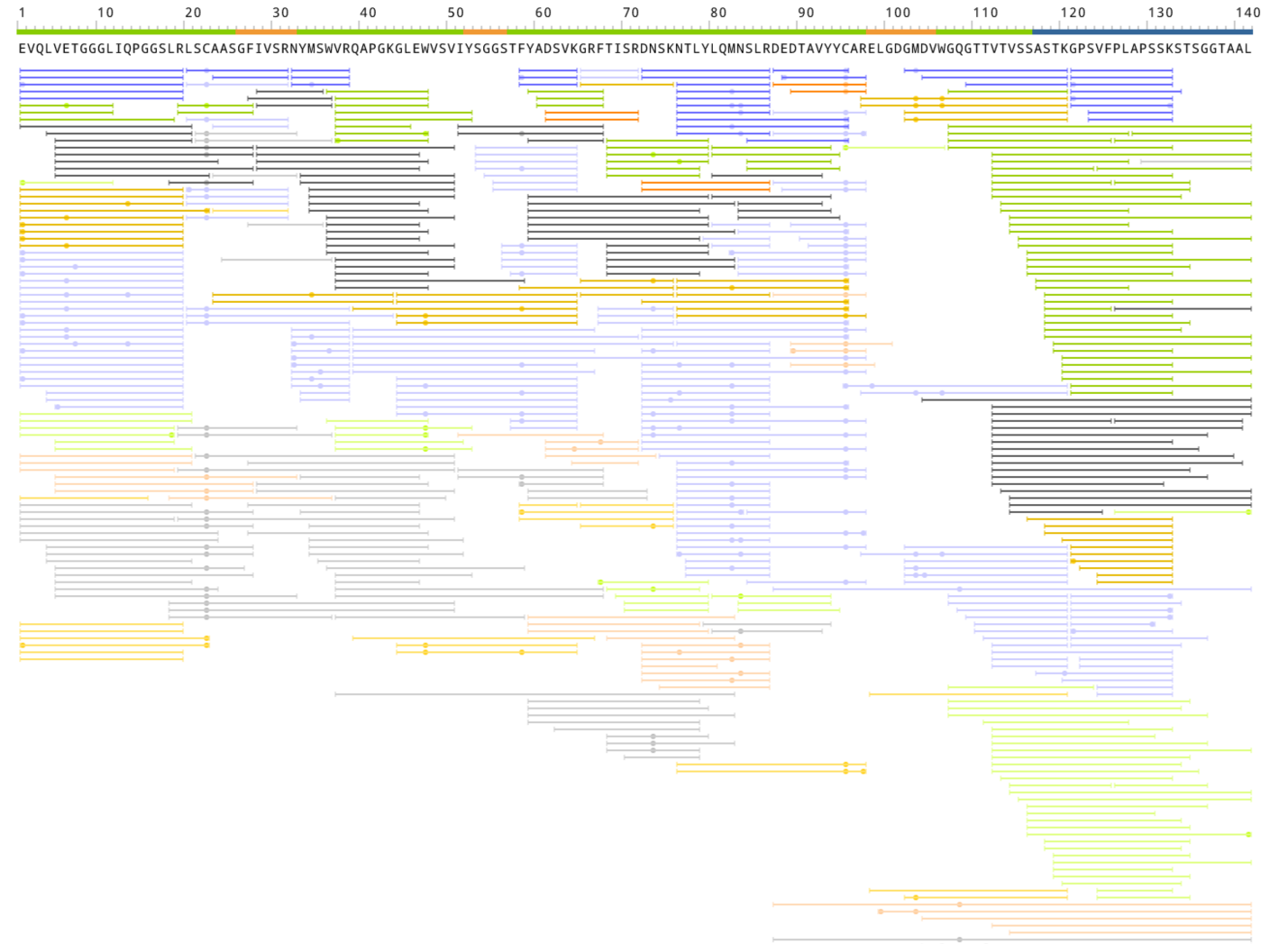

**>heavy-c06.ARVGTTGYDLYGMDV.denovo1_IGHV3-13*04**

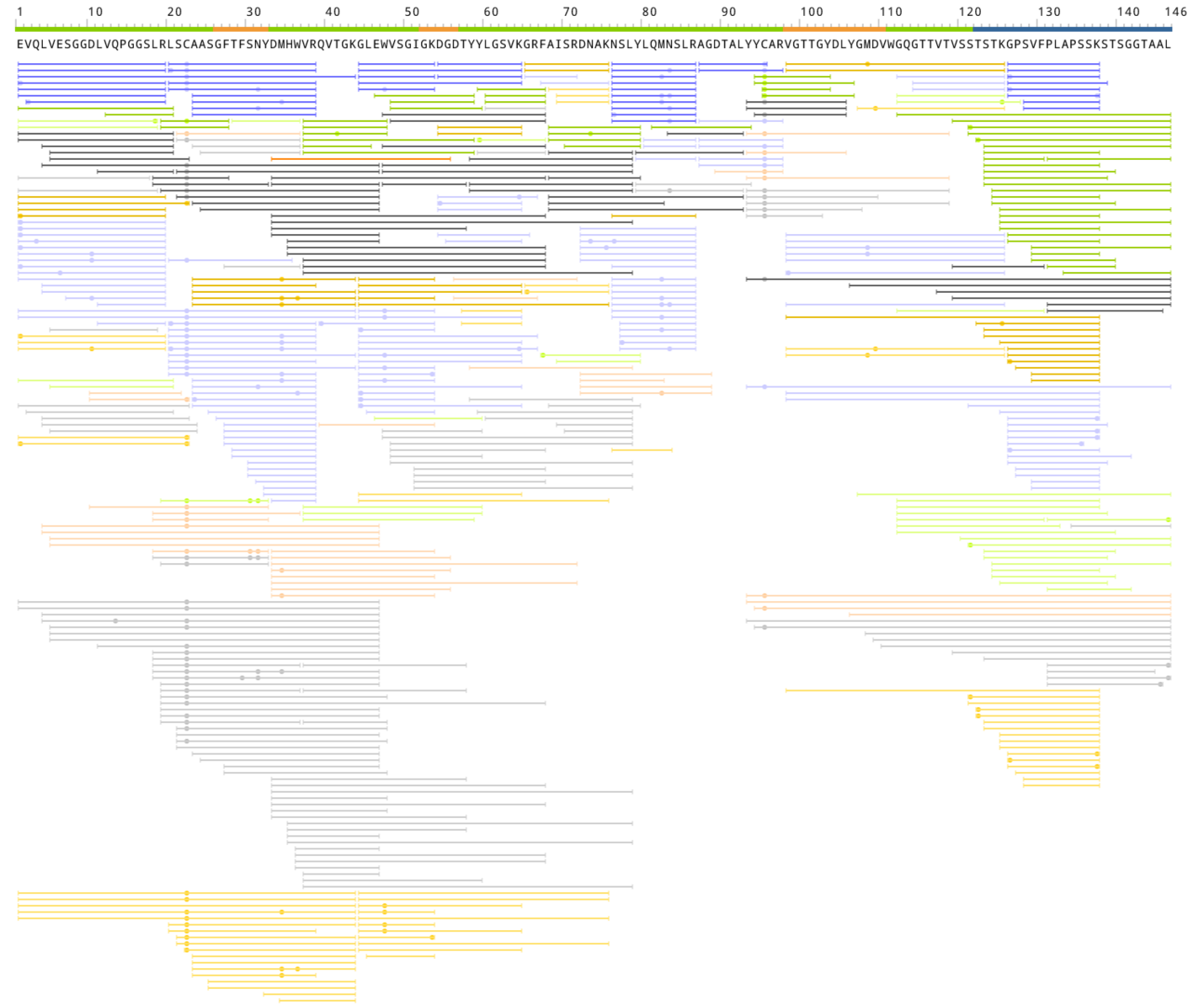

**>heavy-AKTQWEHEGSDFDY.denovo4_IGHV3-23*03**

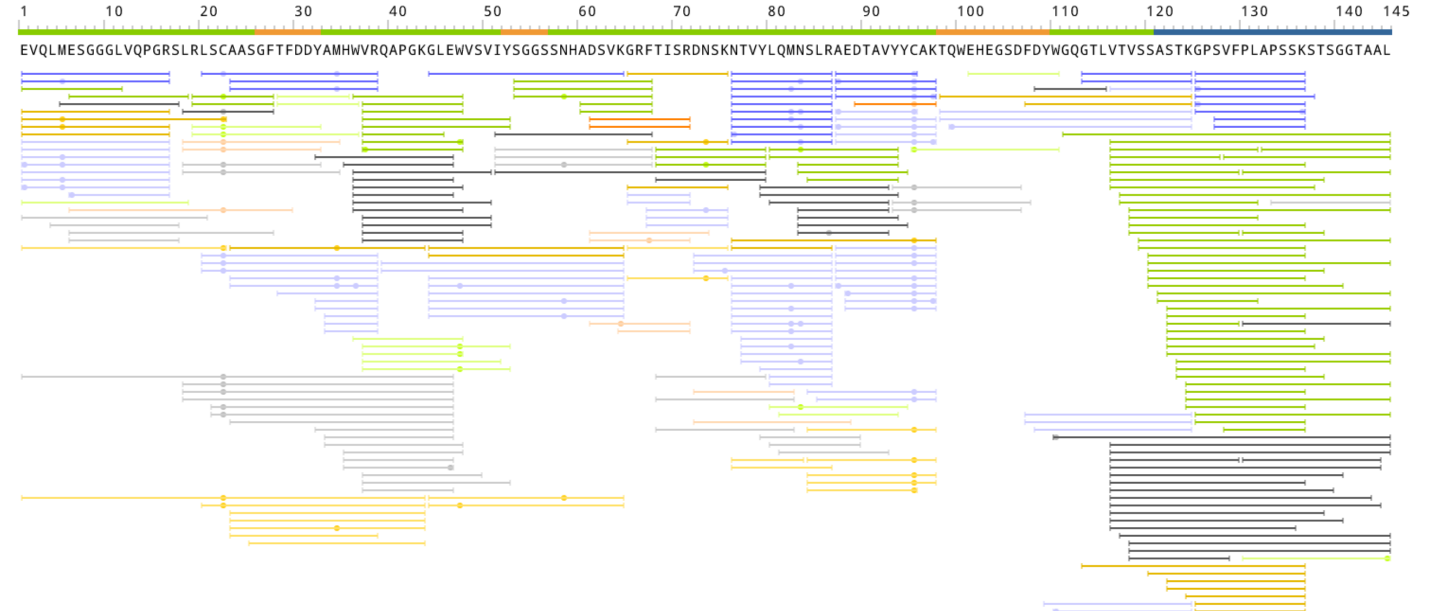

**>heavy-c02.ARYNSGYYALDY.h1RNA_IGHV3-33*08**

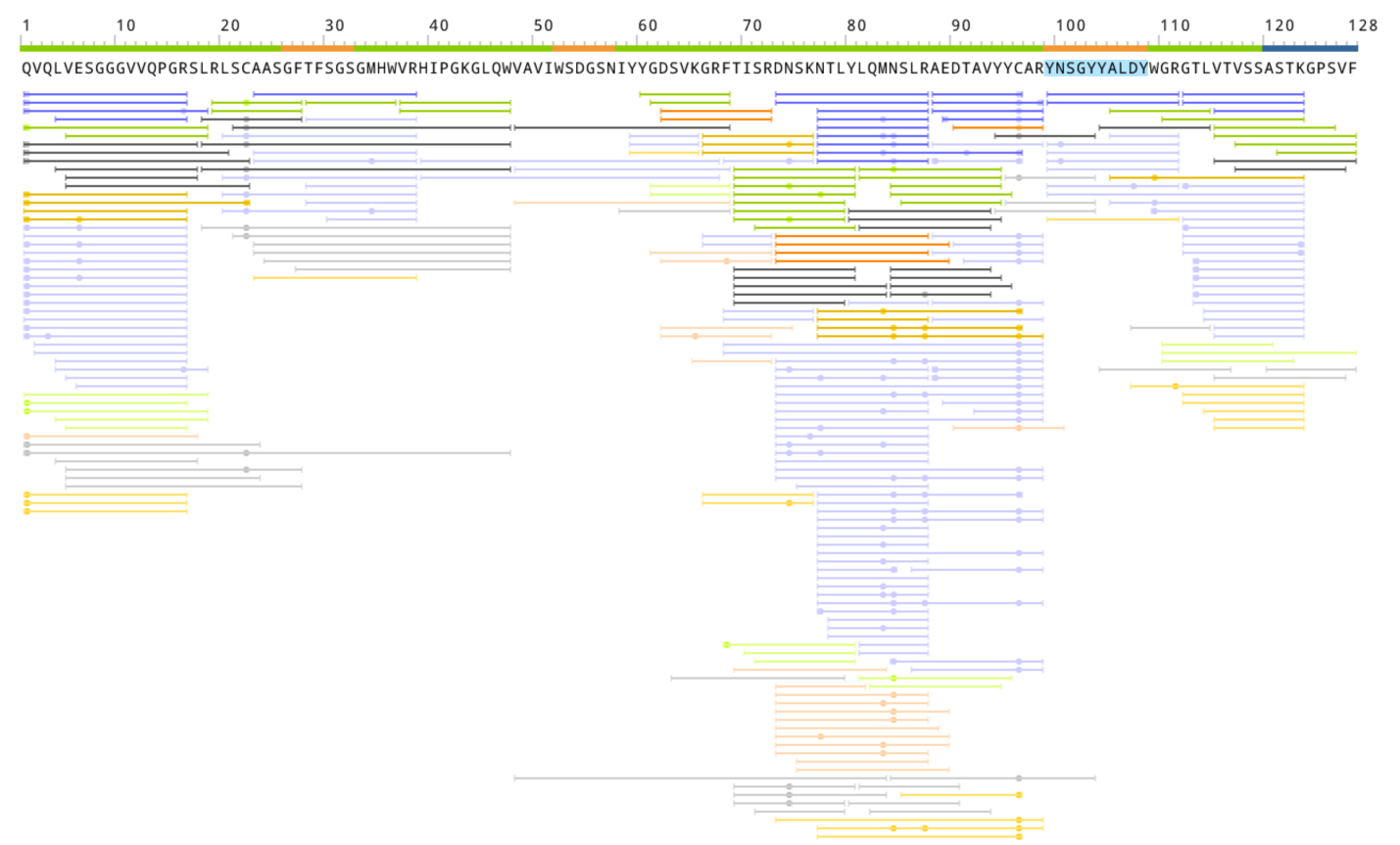

**>heavy.ARKQWLRGAFDI.denovo3_IGHV4-39*09**

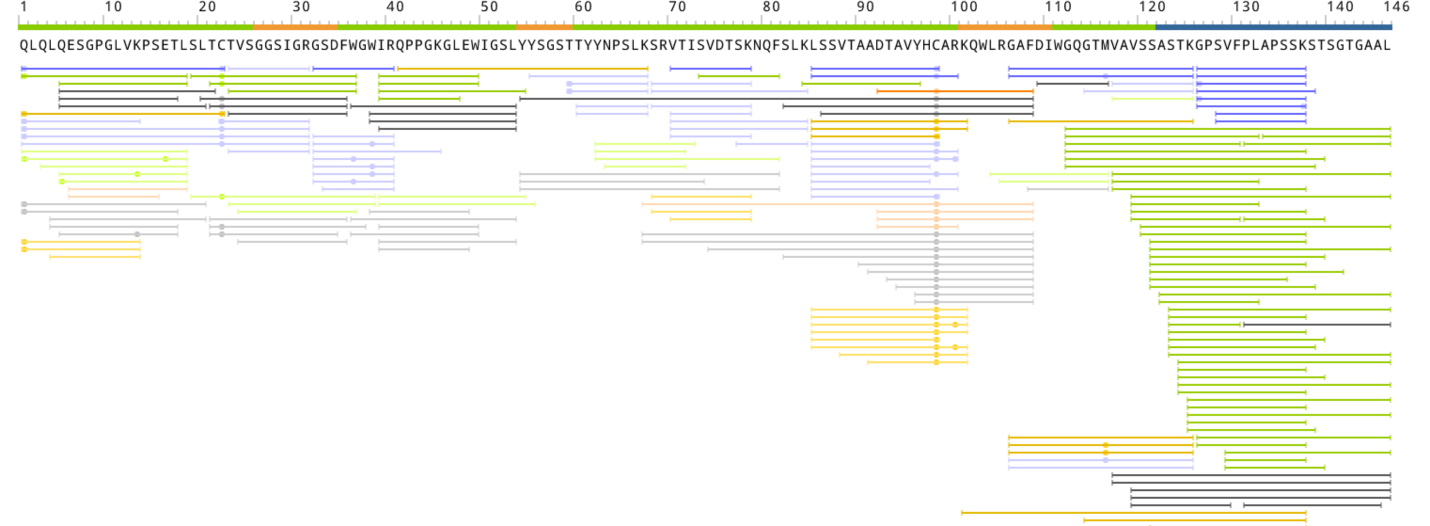

**>kappa-c016.k2.QQLNSYPRT_IGKV1-9*01**

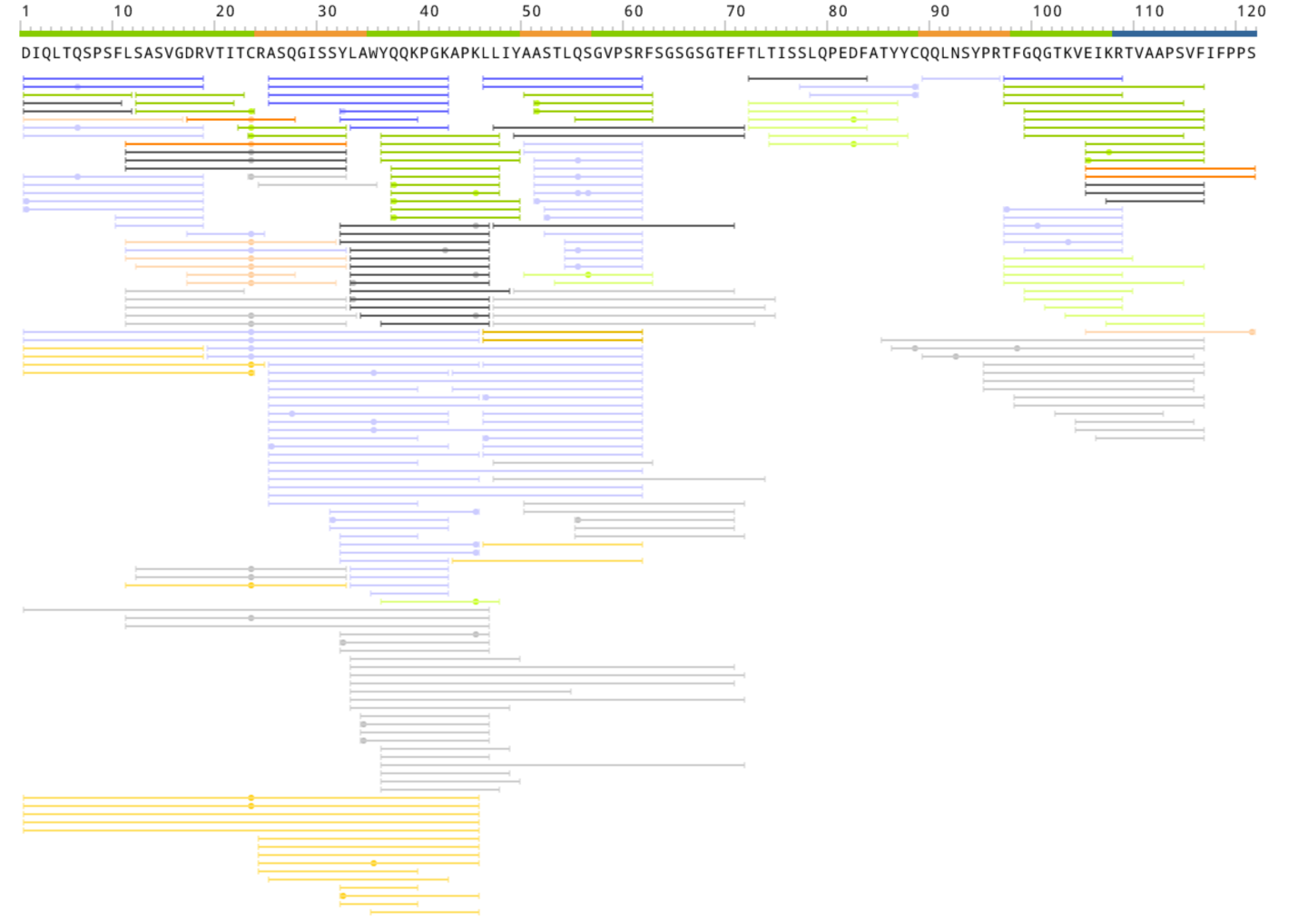

>**kappa-c016.k3.QQLNSYPRGYT_IGKV1-9*01**

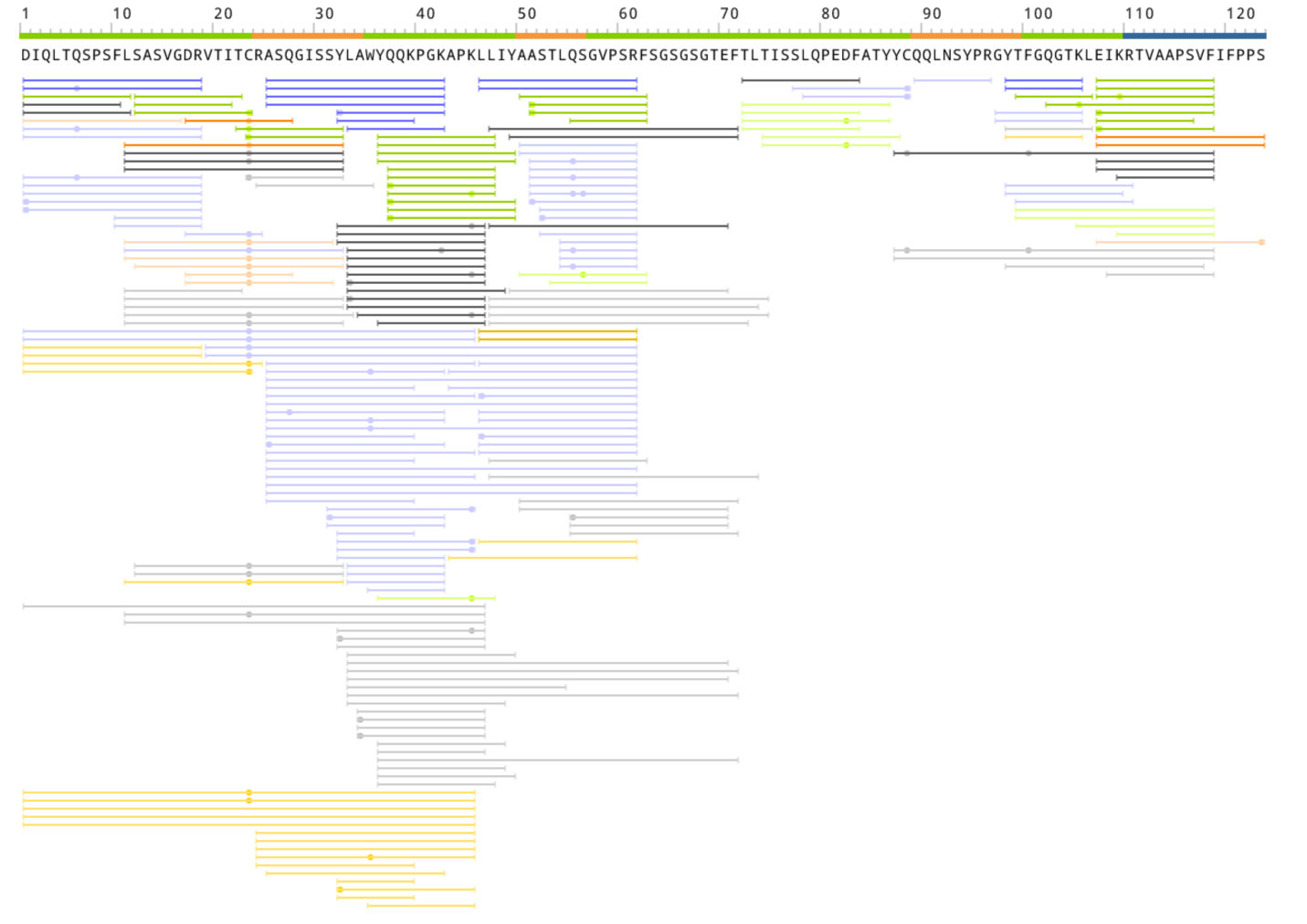

**>lambda-c02.QSGDSGGWV.denovo1_IGLV3-25*03**

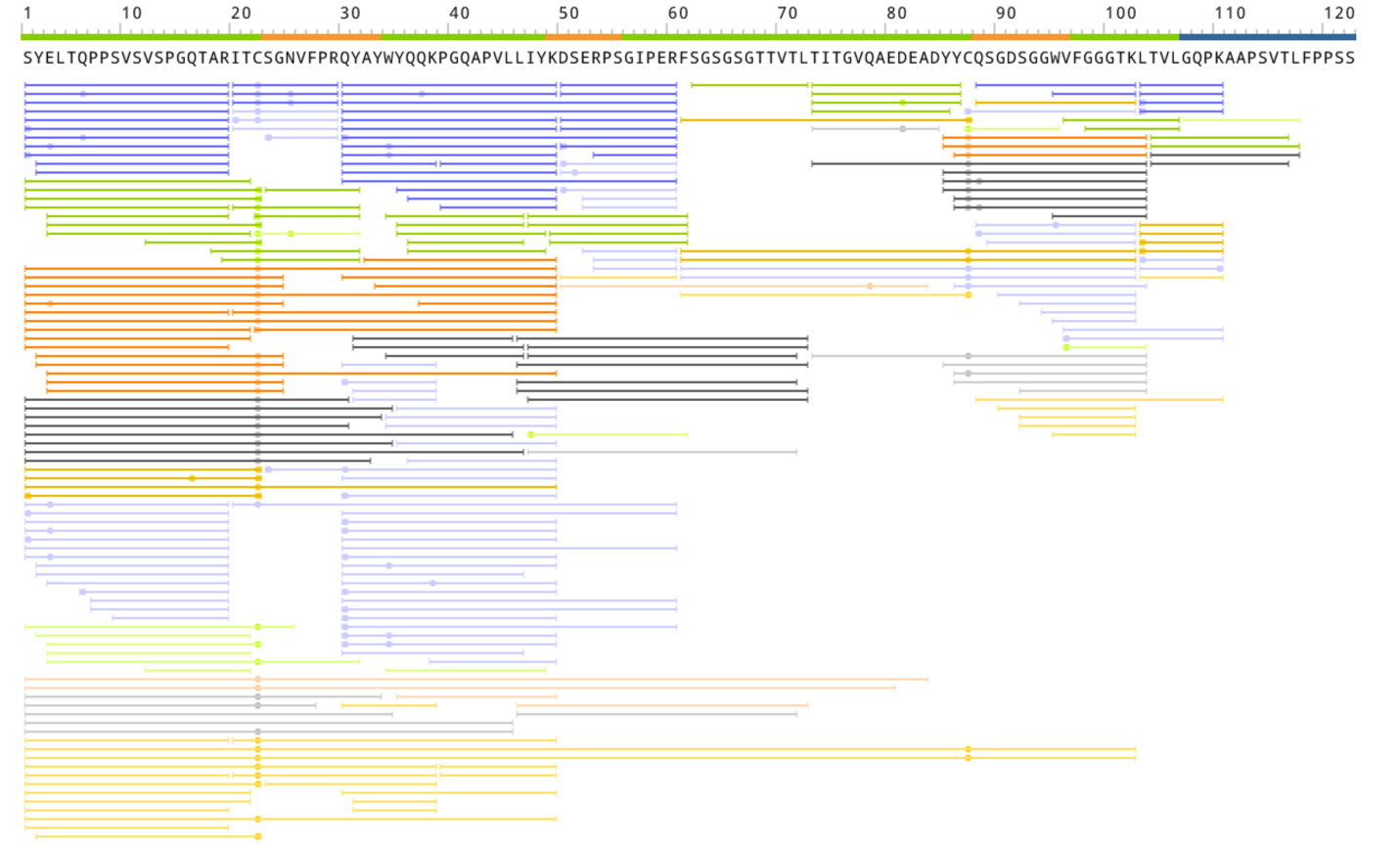

**>lambda-c02.QSGDSGGWV.denovo2_IGLV3-25*03**

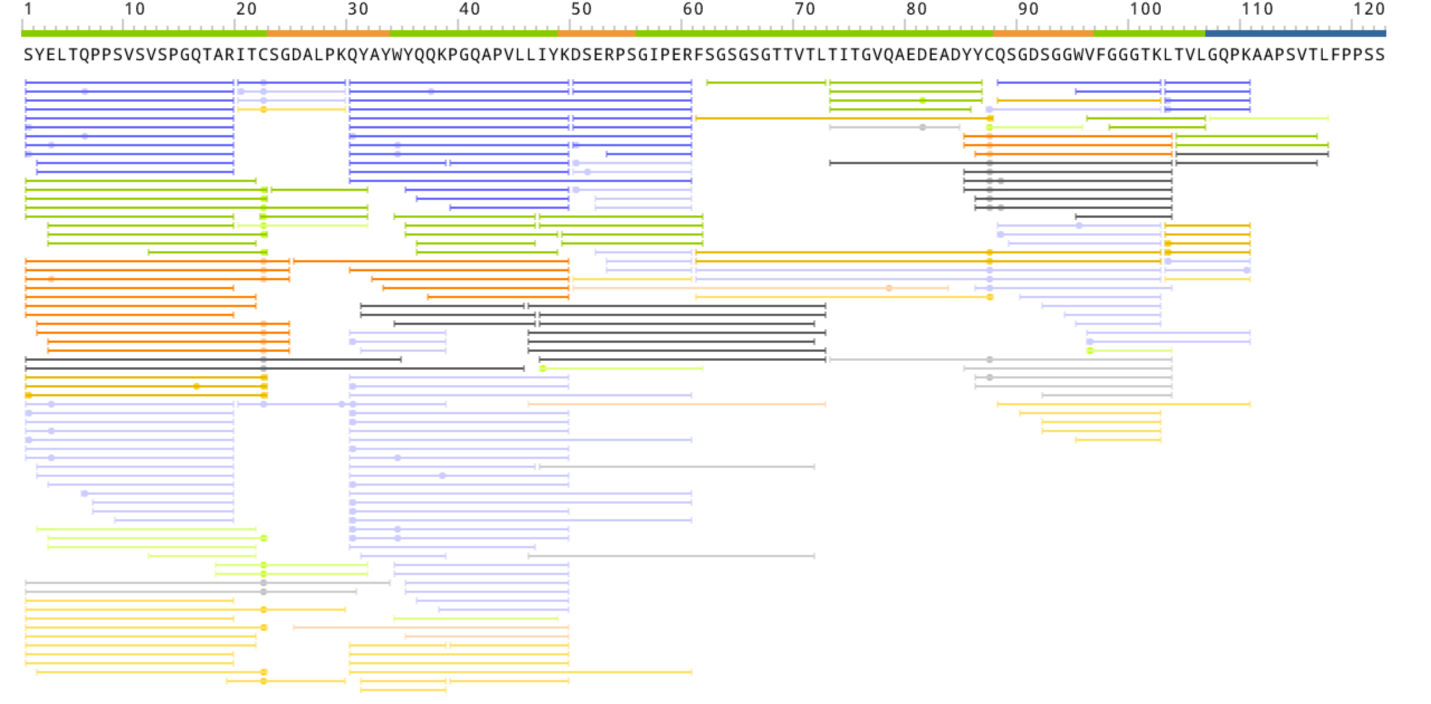

**>lambda-c09.1.QVWDSGTDHSHVV.denovo_IGLV3-21*02**

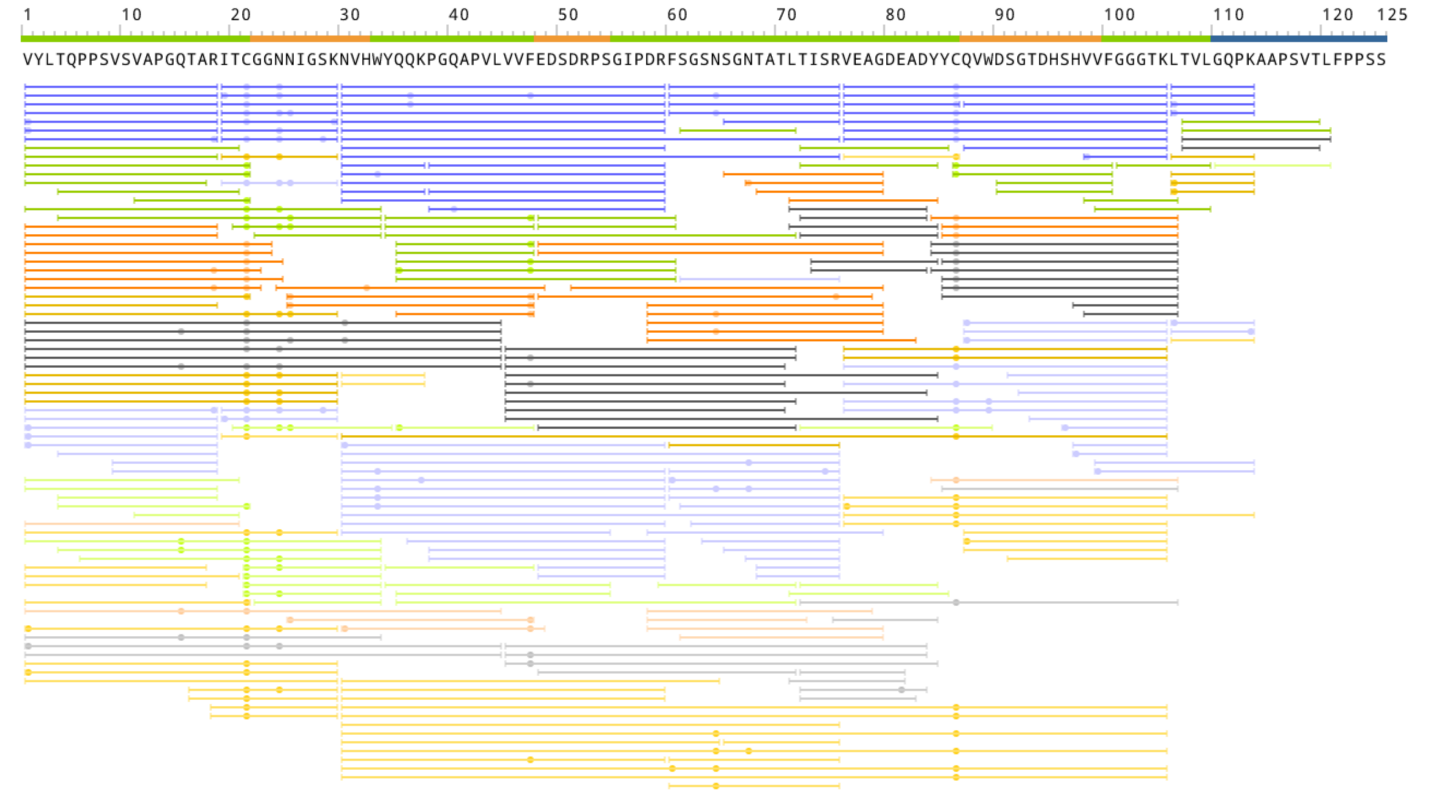

**>lambda-c09.3.QVWDSGTDHSHVV.denovo_IGLV3-21*02**

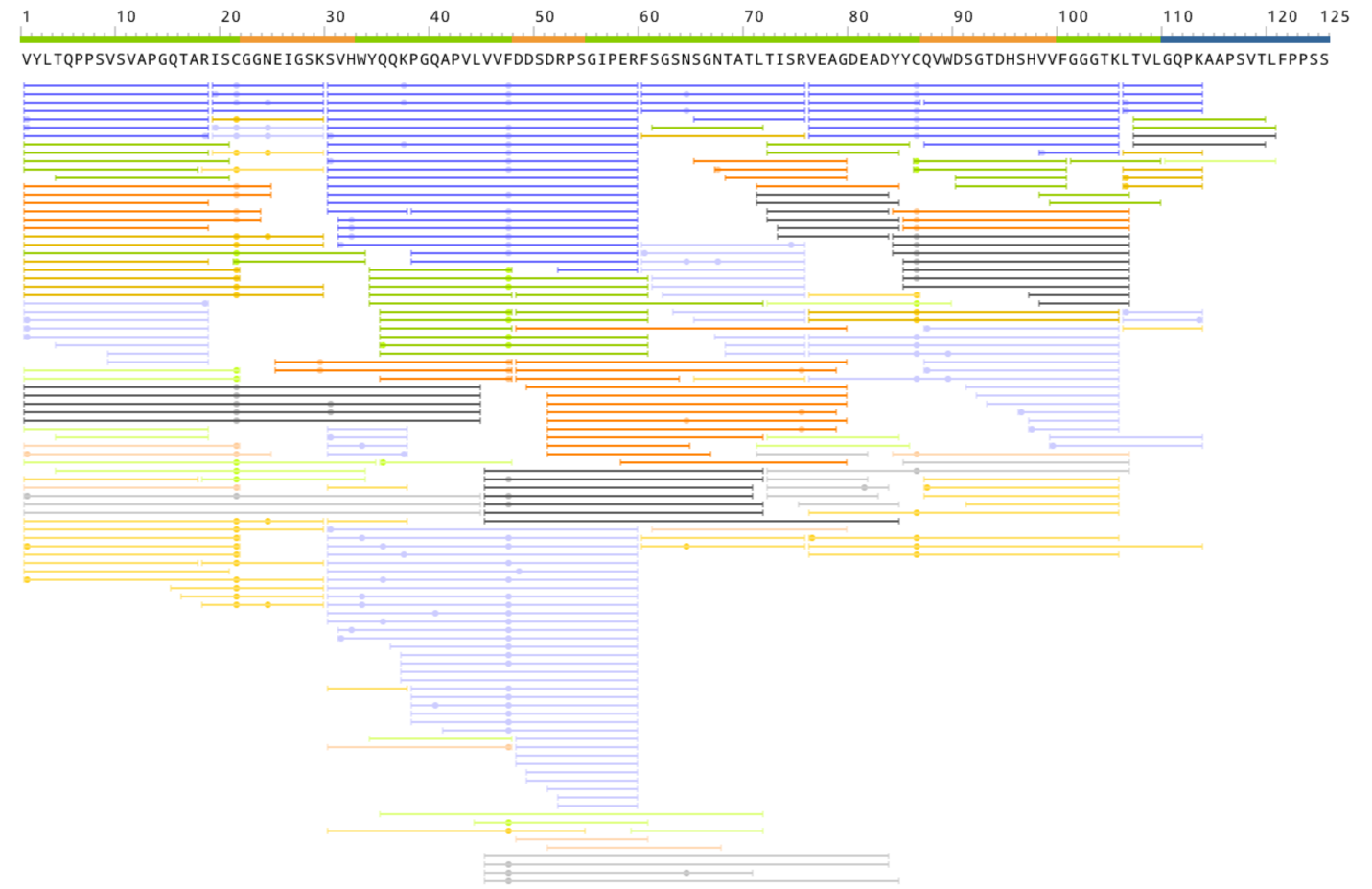

**>lambda-c04.l1.QVWDRSSDHPVV_IGLV3-21*02**

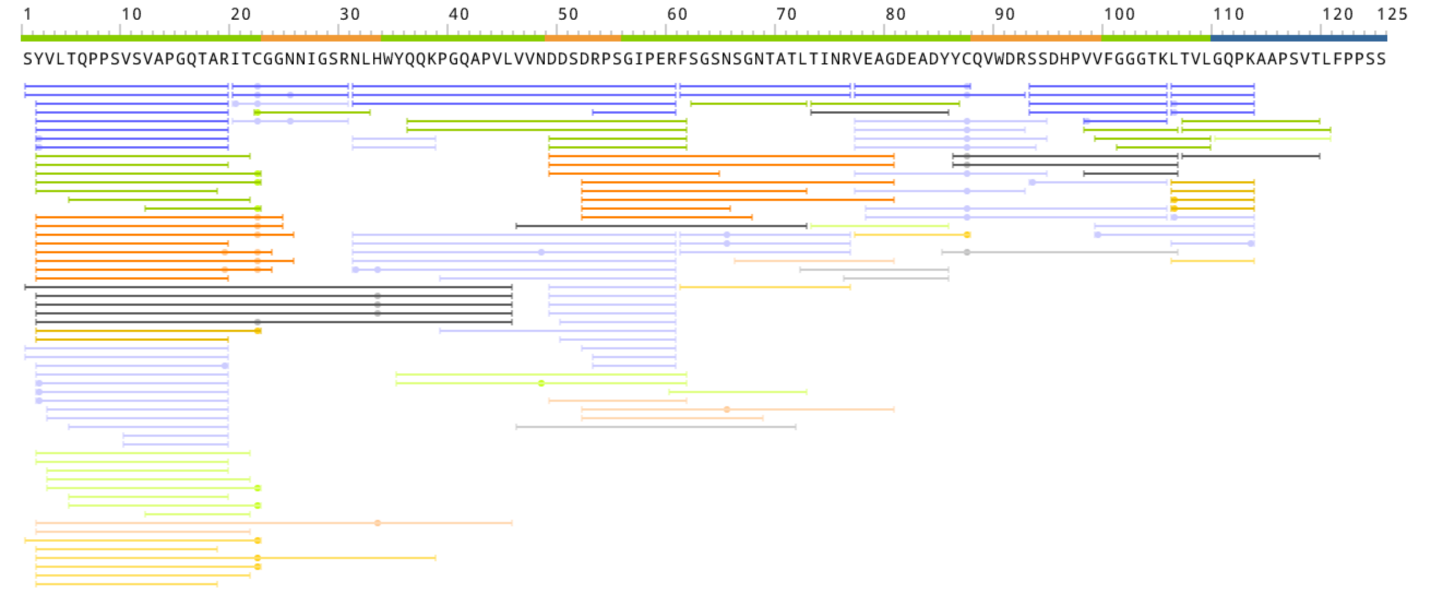

**>lambda-c04.l2.QVWDSSSDVV_IGLV3-21*02**

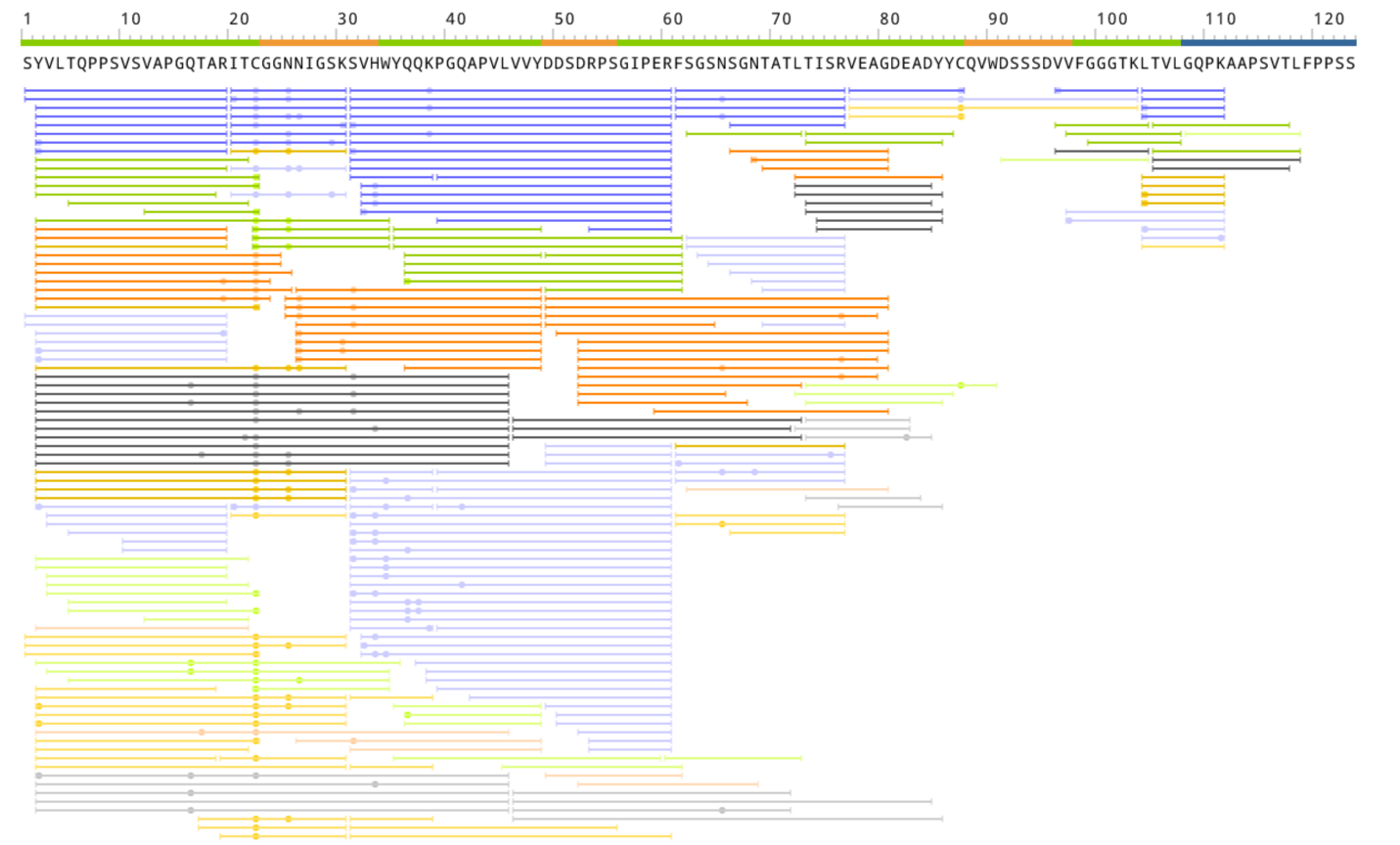
