## Supplementary material for "*de Novo* Sequencing of Antibodies for Identification of Neutralizing Antibodies in Human Plasma Post SARS-CoV-2 Vaccination": SI Appendix

**This PDF file includes:**

Tables S1 to S3

Figures S1 to S2

Supporting Materials and Methods

SI References

**Other supporting materials for this manuscript include the following:**

Datasets S1 to S3

**Table S1: Samples description**

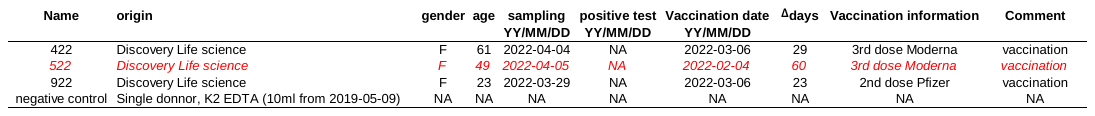

**Table S2: Peptides detected under non-reduced conditions.**

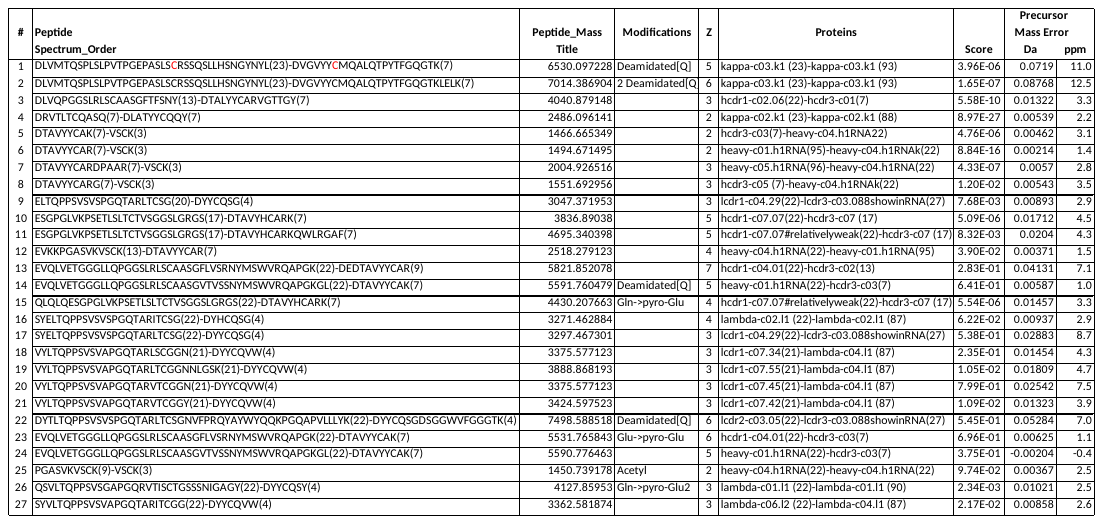

**Table S3: Contigs from Figure 3B**

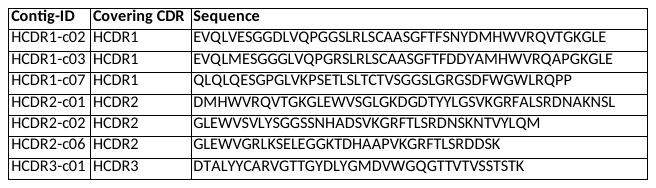

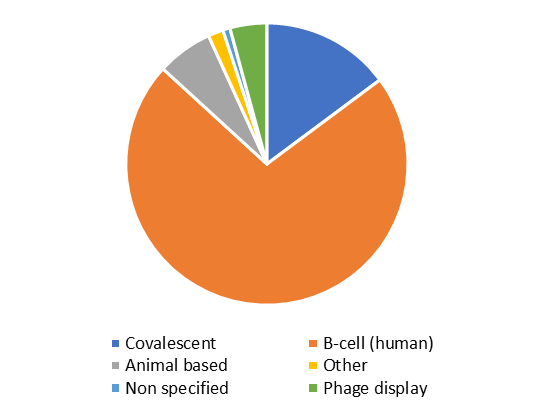

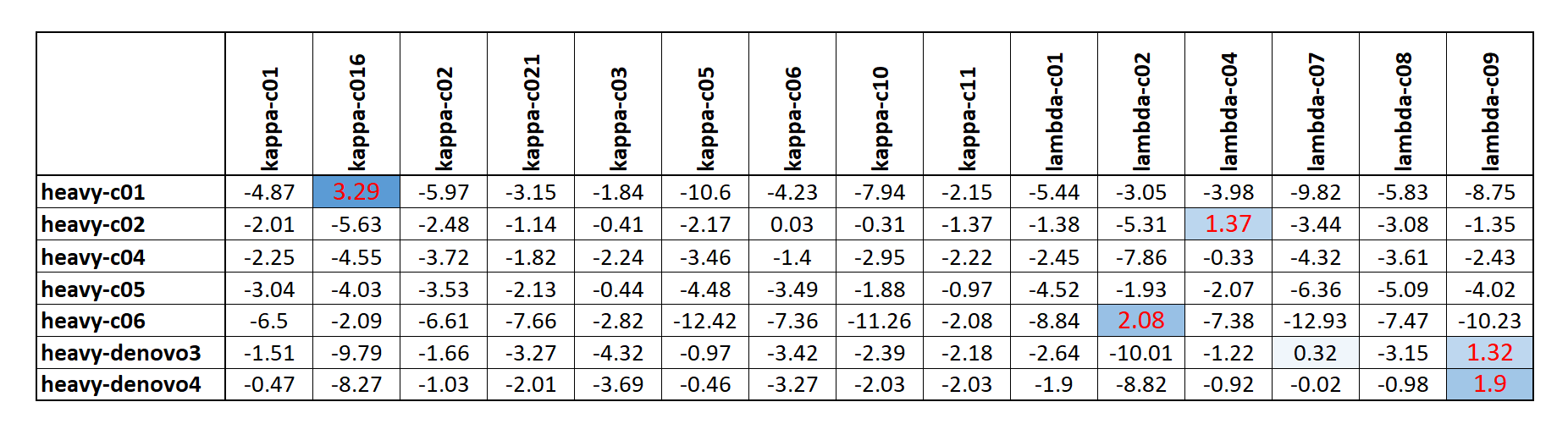

B

A

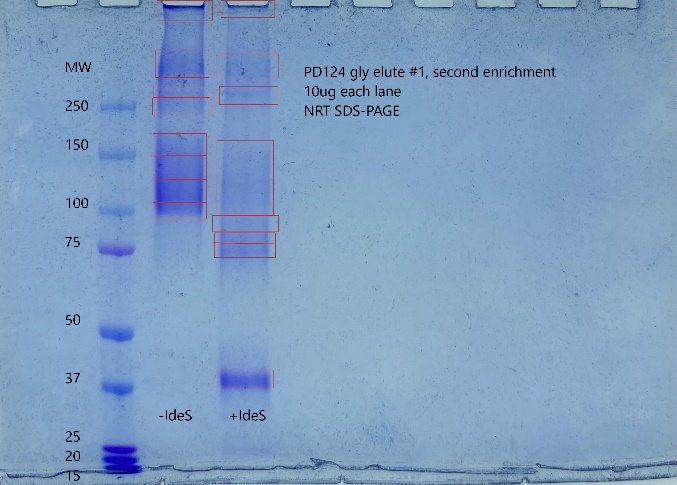

C

**Figure S1**: Additional experiments associated with the polyclonal characterization, the germline, and CoV-AbDab database analysis. **Fig. S1A** illustrates the natural polyclonal separation using "NRT" separation (with and without IdeS). **Fig S1B** shows the best-selected candidates' heavy and light chain pairing using the two separation methods (NRT and Native gel). **Fig. S1C** shows a method breakdown on 12016 sequences extracted from CoV-AbDab.

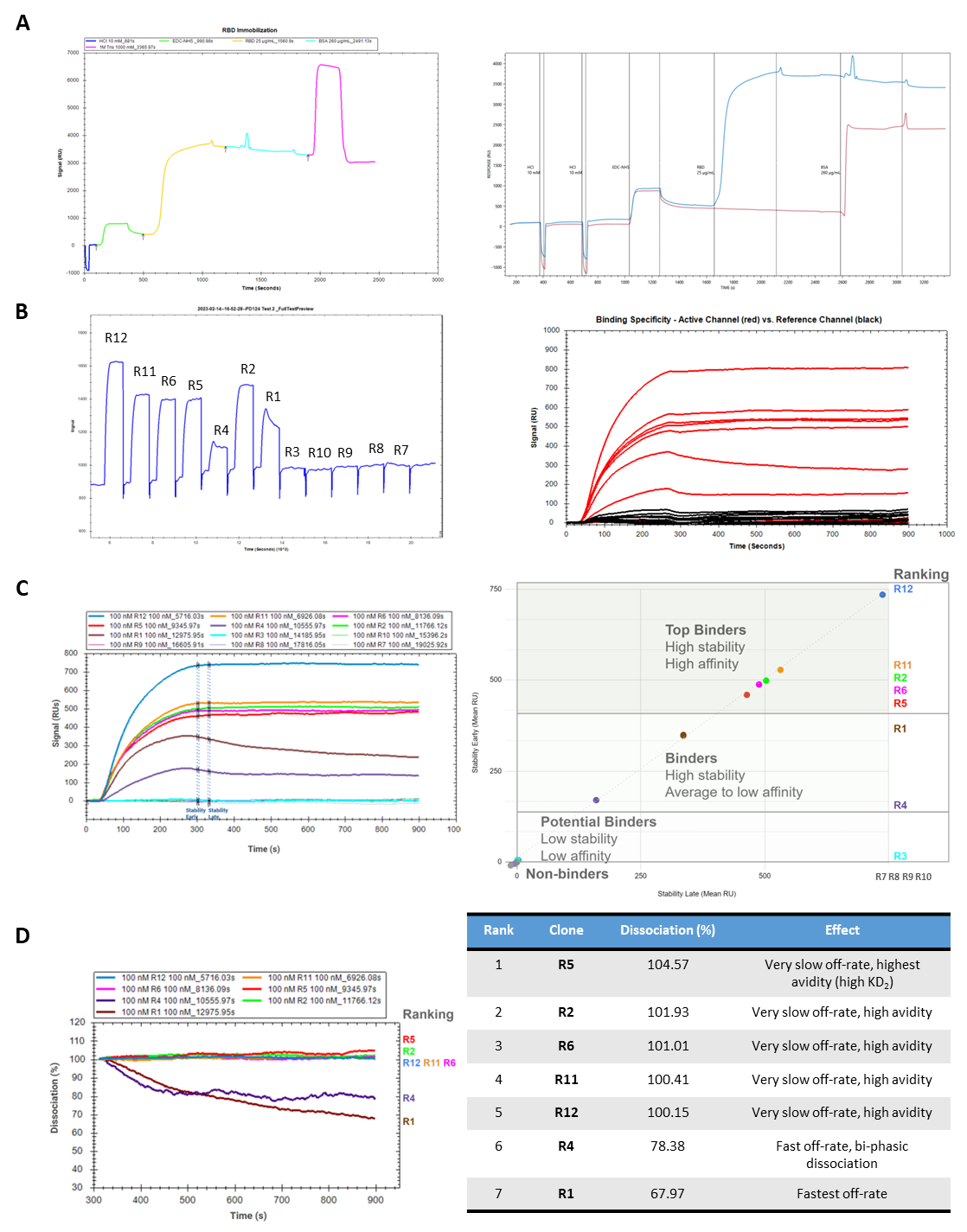

**Figure S2.** Affinity Screen by SPR. **Fig. S2A** Optimized immobilization conditions for RBD on High-Capacity Carboxyl Sensors. Representative sensorgram shows injection sequences for the ligand in Active Channel. Comparative traces (right) for immobilization on the 2-Channel OpenSPR instrument, with the blue trace representing RBD immobilization in the active channel and the red trace representing BSA immobilization in the Reference Channel. **Fig. S2B** Corrected sensorgram of the entire antibody injection sequence with regeneration. Antibody analytes were injected at a fixed concentration of 100 nM in sequence, with a single regeneration cycle of 10 mM Glycine-HCl after each analyte injection. All clones showed >90% regeneration efficacy, as demonstrated by the signal's return to the baseline matching the pre-analyte level. Overlay of analyte responses in the Active Channel with RBD immobilized (red traces) vs. Reference Channel with BSA immobilized (black traces). Binding responses are specific, with a low non-specific binding and bulk shift. **Fig. S2C** A complex stability screen assessed the affinity of antibody clones R1-R12 for immobilized RBD. Response levels were measured immediately at the end of the analyte injection (stability early) and after 30 sec (stability late). Plotting of the two stability values for each clone identified four groups of binders. **Fig. S2D** A dissociation rate, off-rate (kd) screen was performed on clones with validated binding (Top binders and Binders). We normalized the traces for levels of individual binding responses, and percent dissociation was measured for 10 min at 24° C.

**Supporting Materials and Methods**

**Sample collection:** Three samples were collected by Discovery Life Sciences from 3 different healthy donors, with their consent. We are presenting the data from a single donor (named "522"). The patient is a Caucasian female, 49 years old (*SI Appendix* Table S1 for more details). PBMC from blood was collected using BD Vacutainer CPT™ Cell Preparation tube with Sodium Heparin according to supplier’s instructions and stored in RNAprotect® Cell Reagent (Qiagen). Samples for proteomics were collected as follows: blood was centrifuged at 2000 x g 15min 4°C pipette, 50% of the sample volume was transferred into a Matrix cryo vial.

**RNA Extraction**: PBMCs were lysed and homogenized using a QIA shredder (Qiagen) before RNA extraction following the supplier's recommendation. Two 1ml cryovials of human PBMCs from the same patient were used.The PBMC and RNAprotect Cell Reagent mix was centrifuged at 5000 x g for 5 mins to isolate the cell pellet. 600 µl of Buffer RLT containing 6 µl of beta-mercaptoethanol was added to the cell pellet and vortexed until completely dissolved. The lysate was transferred to a QIAshredder spin column and centrifuged for 2 mins at 19,500 x g. The flow-through from the QIAshredder was transferred to a new tube, and the homogenized lysate was combined with an equal volume of 70% ethanol and proceeded to RNA extraction. Following the manufacturer's instructions, the total RNA from PBMCs was isolated using the RNeasy Mini Kit (Qiagen). Extracted RNA concentration was quantified using the RNA BR assay for Qubit fluorometry (Thermo Fisher Scientific), and RNA integrity was determined using the Agilent TapeStation 4150 (Agilent Technologies), which generated an RNA Integrity Number (RIN) of 9.6.

**cDNA synthesis, BCR Amplification, and Sequencing Library Generation:** The SMARTer Human BCR IgG IgM H/K/L Profiling Kit (TakaraBio) was used to construct libraries for NGS. To generate cDNA, one µg of RNA was input into the first strand cDNA synthesis reaction following instructions from the BCR profiling kit. PCR conditions for BCR amplification and sequencing library generation were followed according to the kit user manual using the provided IgG, IgK, and IgL primers, except increasing the recommended cycles for PCR2 from 16 to 18. The concentration of each library was quantified using the 1X dsDNA HS assay for Qubit fluorometry (Thermo Fisher Scientific), and the library sizes were visualized using the D1000 ScreenTape on the Agilent TapeStation 4150 (Agilent Technologies). The expected library sizes were approximately 640bp for the light chains and 690bp for the heavy chains. Peaks around 350bp are indicative of adaptor dimers, and libraries containing these peaks were subject to a size selection process using the NEBNext Sample Purification Beads (0.65X beads to sample volume) following the SPRIselect User Guide.

**Next-Generation Sequencing:** The final products were re-quantified using the 1X dsDNA HS assay (Thermo Fisher Scientific), and the amplicon sizes were visualized using the D1000 ScreenTape on the Agilent TapeStation 4150 (Agilent Technologies). The final quantified libraries were normalized and pooled to a 4nM library, which was used to prepare an 8 pM loading library. To ensure complete denaturation of the libraries and enable efficient binding to flow cell, the libraries were heat-shocked at 96°C for 2 mins and immediately transferred to water slurry for 5 mins. Sequencing was conducted on an Illumina MiSeq platform using the reagent kit v3 with 600 cycle paired-end reads with a 30% PhiX v3 spike-in.

**IgG enriched against the antigen**: Total IgG was from 3 mL human serum from a patient vaccinated against SARS-CoV-2. Briefly, 3 mL of settled protein G agarose resin (Genscript) in a 20 mL gravity flow Biorad column was equilibrated by washing twice with 15 mL 10 mM phosphate buffered saline (PBS). To remove debris, serum was prepared for enrichment by centrifuging at 23,000 x g for 10 min at 4°C. The 3 mL serum was combined with 9 mL PBS, then filtered. Filtered serum was passed over protein G agarose resin using gravity flow. Protein G resin was washed with PBS and then eluted using glycine buffer 0.1M

pH 2.5. Eluted IgG was then concentrated, and the buffer was exchanged into PBS using a 30 kDa Amicon filter (Sigma-Aldrich). The total amount of IgG captured was found to be 22.2 mg.

Antigen enrichment was performed to enrich anti-RBD antibodies by coupling SARS-CoV-2 spike protein receptor binding domain, RBD, (ExonBio) to streptavidin-coated agarose beads (Sigma-Aldrich) for 1 hour at 4 °C with tumbling. Biotinylation of RBD was performed using 20 molar times excess of photocleavable biotin, PC biotin-PEG3-NHS ester (Sigma-Aldrich) reconstituted in acetonitrile, incubated at room temperature for 1 hr, followed by a buffer exchange into 10 PBS using a 3 kDa Amicon filter (Sigma-Aldrich) to remove excess biotin. Following the coupling of biotinylated RBD to streptavidin agarose resin, three washes were performed using 0.4 mL 10 PBS to remove any uncoupled RBD, then 20 mg total IgG was added and incubated for 1 hour at 4°C with tumbling. Non-specific binders were removed by washing twice with 0.4 mL PBS, followed by a wash with 0.4 mL CHAPS 0.5% in PBS, then six washes with 0.4 mL PBS. Anti-RBD antibodies were eluted twice using 0.4 mL glycine pH 2.5, incubated for 5 min at room temperature and combined, then neutralized using 0.2 mL 1 M tris pH 8, resulting in 111 ug anti-RBD antibodies, which were given an internal sample name of PD124. Beads were washed with 100 uL PBS, then 400 uL PBS to bring pH to 7, then 1 hr UV elution was performed in 100 uL PBS using UV Stratalinker 2400 (Stratalinker), with 365 nm bulb to cleave and elute any remaining anti-RBD antibodies. UV elution resulted in an additional 14.2 ug PD124 eluted.

**In-solution Digestion:** An in-solution digestion was performed on 25 ug of the sample PD124 by concentrating the sample to 100 µL using centrifugation under low pressure (i.e., Speedvac), followed by reduction using dithiothreitol (DTT) at a final concentration of 30 mM at 95°C for 15 minutes. The sample was then split into two: 5/7th was alkylated with Iodoacetamide (IAA) at a final concentration of 50 mM at room temperature in dark conditions for 30 minutes, and 2/7th was treated with 2-bromoethylamine hydrobromide (BEA) at a final concentration of 0.5 M for 4 hours at 25°C with shaking. The sample treated with iodoacetamide (IAA) was then precipitated by adding acetone to a final concentration of 75% of the sample and incubating for 1 hour at -20°C, then centrifuging at 23,000 x g for 10 mins at 4°C. Acetone was decanted, and the pellet was dried under low pressure, followed by reconstitution with 10 µL 4 M urea, which was incubated at 37°C for 10 min with shaking to reconstitute the pellet. The sample was then split across five tubes, and 30 µL of 50 mM ammonium bicarbonate was added to 4 for digestion with Trypsin, LysC, AspN, and Chymotrypsin at a 1:20 ratio of protease: protein. These samples were left to digest overnight in a 37°C incubator. For the fifth digestion, 30 µL of HPLC grade water and 2 µL of 1N HCl and pepsin were added at a 1:20 ratio of protease: protein. Pepsin digestion occurred at 37°C for 15 minutes with shaking, followed by a 3-minute pepsin inactivation at 95°C. Digests were then dried under low pressure.

BEA-treated samples were precipitated by adding trichloroacetic acid to a final volume of 20% and then incubating overnight at 4°C. The following day, samples were pelleted with centrifugation at 14,000 x g for 30 minutes at 4°C, then pellets were washed twice with 500 µL 80% acetone with centrifugation at 14,000 x g for 10 min for each wash step. Pellets were dried under low pressure and then reconstituted in 4 µL 4 M urea at 37°C for 10 min with shaking, then split across two tubes and digested in ammonium bicarbonate at a final concentration of 30 mM with Trypsin and LysC at a ratio of 1:20 protease: protein. Digestions were incubated at 37°C overnight. All proteases used for PD124 digests were obtained from Promega.

Digested samples were dried to completion in a centrifuge under low pressure, then resuspended in 40 µL 0.1% formic acid. 2.5 ug was loaded on Evotips according to the manufacturer's instructions for each digestion.

**Native gel:** PD124 was separated on the gel using Biorad precast 7.5% polyacrylamide gels with and without IdeS treatment. For IdeS treatment, 50 ug of PD124 was incubated at 37°C with 50 units of IdeS from Promega for 1 hour, then dried under low pressure to reduce volume to 30 µL. Nativepage 4x buffer (Thermo Fisher Scientific) was added in a 4:1 ratio. For the undigested sample, only 25 ug was loaded on gel after adding NativePage 4x buffer in a 4:1 ratio. Running buffer was made by diluting 10X stock Tris/Glycine Buffer (Biorad) to 1X using Milli-Q water. The gel was run using a Biorad power bank set to 130V for 180 min, then stained with Coomassie brilliant blue (Biorad) for 30 min and destained with Biorad Destain overnight.

Native in-gel digestion: Bands were excised and washed with water following gel separation.

The gel was cut into 12 bands (Figure 3A). Trypsin was used to digest each band using a standard in-gel digestion protocol (1).

**NRT gel (Figure S1A):** PD124 was separated using a modified non-reducing SDS-PAGE for heavy–light chain pairing where the sample was not heated before electrophoresis. The separation was performed using 10 µg per lane of sample in its native form or following digestion with the protease IdeS. Ides digestion was performed at 37°C for 1 h with shaking in the sample buffer. Samples were loaded on precast 7.5% Mini-PROTEAN TGX gels from Biorad using lithium dodecyl sulfate (LDS) sample loading buffer at a 4:1 ratio. Gels were run using premade stock solutions of Tris/Glycine/SDS Buffer (Biorad) and the PowerPac Basic power supply at 150V for 60 min (Biorad). Trypsin was used to digest each band using a standard in-gel digestion protocol(1).

**Digestion under non-reducing condition:** For non-reduction digestion analysis, the sample preparation consisted of using 20 µg of the polyclonal antibody dried completely under low pressure, then reconstituted in 20 µL 8 M urea in 100 mM tris buffer, adjusted to pH 6.5, N-ethylmaleimide (NEM) was added to a final concentration of 2 mM and incubated at 37°C for 2 hours with shaking. The endoprotease, LysC was added at a molar ratio of 1:20 (protease to protein ratio) and incubated at 37°C for overnight digestion, followed by 4-hour digestion with AspN at a ratio of 1:20 (protease to protein ratio) at 37°C. After protease digestion, the sample was dried under low pressure and reconstituted in 40 µL of 0.1% formic acid. Then, 2.5 µL of the digested samples were loaded onto Evotips according to the manufacturer's instructions.

**Isobaric resolution:** Three selected samples were analyzed in EThCD mode to resolve potential isobaric ambiguity (such as isobaric amino acid I/L). Those samples are: the previous trypsin and pepsin digest were used as is, and the pepsin one was, in addition, labeled at the C-terminal end using an approach similar to Krusemark et al. (2). However, we used arginine methyl ester dihydrochloride (Sigma-Aldrich) as the primary amine reagent.

**Mass spectrometry analysis:**

***HCD mode:*** Samples from in-solution, ingel, and non-reduce digestion were all run on Orbitrap 240 Exploris (Thermo Fisher Scientific) using 30 samples per day, 44 min method on a 15cm PepSep column. Data was acquired at 60,000 resolution for 400-2000 m/z precursors mass range with standard AGC target and maximum injection time set to auto. An intensity threshold of 2.5e4 was applied, and charge states 2-8 were included. Dynamic exclusion was set to 15s. MS/MS fragmentation was induced with fixed 30% HCD, detected by the Orbitrap with 7,500 resolution with a maximum injection time of 50ms.

***EThcD mode:*** The sample for isobaric resolution were analyzed using an Orbitrap Eclipse Tribid Mass spectrometer instrument (Thermo Fisher Scientific) connected to an Evosep operated in a 30 sample per day, 44min method on a 15cm PepSep column. Data was acquired at 60,000 resolution for a m/z range of 400-2000 m/z precursors mass range with standard AGC target and maximum injection time set to auto. An intensity threshold of 2.5e4 was applied, and charge states 2-8 were included. Dynamic exclusion was set to 15s. MS/MS fragmentation was induced in ETD, with SA collision Energy type set at EthCD, with fixed energy at 55. MSMS spectra were acquired at 7,500 resolution with a maximum injection time of 50ms.

**ELISA**

***Titer evaluation from plasma:*** For the ELISA with plasma, 0.1ug recombinant RBD (ExonBio) was immobilized on Maxisorp 96-well plates (Thermo Fischer Scientific) in sodium carbonate-bicarbonate buffer (Sigma-Aldrich). The wells were washed with PBS-T and blocked for 1 hour at room temperature with SuperBlock buffer (Thermo Fischer Scientific). Plasma dilutions were 1:30, 1:90, 1:270, 1:810, 1:2430, 1:7290, 1:21870 with Stabilzyme diluent. After washing, the wells were incubated with serially diluted plasma in duplicate for 1 hour. The wells were washed and incubated with goat anti-human secondary antibody in a 1:5000 dilution (Sigma-Aldrich). The wells were washed, and the TMB substrate (Thermo Fischer Scientific) was added. The reaction was quenched with a 0.2N sulfuric acid (Sigma-Aldrich). The plates were read in a spectrophotometer (Biotek Synergy LX) at 450nm. The average OD450 for each sample was plotted on a log scale.

***Polyclonal profiling and recombinant forms testing:*** 0.2ug recombinant RBD (Exon Bio) was immobilized on Maxisorp 96-well plates (Thermo Fischer Scientific) in sodium carbonate-bicarbonate buffer for those ELISA assays. The wells were washed with PBS-T and blocked with 200ul SuperBlock buffer (Thermo Fischer Scientific).). After washing, the wells were incubated in duplicate with serially diluted recombinant antibodies (Thermo Fischer Scientific). Antibody dilutions ranged from 100nM to 0.03nM. The wells were washed as above and then incubated with goat anti-human secondary antibody in a 1:5000 dilution. The wells were washed as above, and then the TMB substrate (Thermo Fischer Scientific) was added to the wells and incubated. The reaction was quenched with a 0.2N sulfuric acid stop solution (Sigma-Aldrich). The plates were read in a spectrophotometer (Biotek Synergy LX) at 450nm. Each sample's average OD at 450nm was plotted on a log-scale graph.

***Surrogate neutralization ELISA (snELISA, in vitro assay):*** For the surrogate neutralization ELISA, 0.2ug recombinant RBD (ExonBio LLC) was immobilized on Maxisorp 96-well plates overnight (2ug/mL in sodium carbonate bicarbonate buffer). The wells were washed with PBS-T and blocked with 200ul Superblock (Thermo Fischer Scientific). After washing, as described above, the wells were incubated with serially diluted recombinant antibodies in triplicate. Antibody dilutions ranged from 100nM to 0.001nM. A standard Sars-CoV2 Spike RBD neutralizing antibody (SAD-S35-100UG from Acro Biosystems) and human IgG from serum (Innovative Research) were used as positive and negative controls, respectively. The wells were washed as above and incubated with biotinylated ACE2 (Sino Biological). After washing as above, the wells were incubated with streptavidin-HRP conjugate (Sigma-Aldrich). The wells were washed as above, and then TMB substrate was added and incubated. The reaction was quenched with a 0.2N sulfuric acid stop solution (Sigma-Aldrich). The plates were read in a spectrophotometer (Biotek Synergy LX) at 450nm. The average OD450 for each sample was plotted on a log-scale graph to produce the neutralization curve.

***In vivo Neutralization assay with pseudovirus assay (Cell-based assay):*** Acro Biosystem performed the assay.

*Cell Culture and Reagents:* HEK293/Human ACE2 Stable Cell Line (Acro Biosystems) was used in this study. The cells were cultured in a DMEM medium supplemented with 10% fetal bovine serum. The complete DMEM medium was prepared by mixing 89% DMEM medium, 10% fetal bovine serum, and 1% penicillin-streptomycin. The HEK293/Human ACE2 Stable Cell Line was cultured in the complete DMEM medium in a CO2 incubator at 37°C with 5% CO2. Serial dilutions of the recombinant antibody samples to test were prepared using the complete DMEM medium. The tested human recombinant antibodies (samples) and positive control (Acro Biosystem) were series diluted at a ratio of 1:4 and were added to 96-well white flat bottom plate wells, each receiving 75 μL of the sample solution. The pseudovirus SARS-CoV-2 Spike (WT) Fluc-GFP Pseudovirus was diluted 125-fold with the complete DMEM medium. The diluted pseudovirus solution (25 μL) was added to each well of the 96-well plate. For the cell control group, 25 μL of the complete DMEM medium was added instead. The reagents in the 96-well plate were gently mixed. The plate was then incubated in a CO2 incubator at 37°C with 5% CO2 for 60 minutes. The concentrations of the samples at this time were recorded for further analysis. Following the incubation, cells were harvested, and cell number and viability were determined. The cell density was adjusted to 5E5 cells per milliliter using the complete DMEM medium. Then, 100 μL of the cell suspension was seeded into each well of the 96-well plate. The plate containing the reagents and cells was gently mixed and then placed back into the CO2 incubator at 37°C with 5% CO2. The plate was incubated for 48 hours. After the incubation, 100 μL of the medium was removed from each well, and the plate was incubated at room temperature for 10 minutes. Next, 100 μL of the detection reagent (Britelite plus Reporter Gene Assay System) was added to each well. The reagent was mixed well with the Mini shaker for 2 minutes.

*Luminescence Measurement:* The luminescence value of each well in the plate was measured using a microplate reader. The detection time for each well was set to 0.1 sec./well.

**Inhibition rate**$\boldsymbol{=}\left( \boldsymbol{1-}\frac{\boldsymbol{X -}\bar{\boldsymbol{CC}}}{\bar{\boldsymbol{VC}}\boldsymbol{-}\bar{\boldsymbol{CC}}} \right)\boldsymbol{\times100\%}$

X: the luminescence value (RLU) of a given well.

CC, cell control, only cells are added.

$\bar{\mathrm{CC}}$ , the mean value of the cell control group.

VC, virus control, only cells, and pseudovirus are added.

$\bar{\mathrm{VC}}$, the mean value of the virus control group.

Analyze the data using GraphPad Prism 8 software (Nonlinear regression). IC50 or relative IC50 is defined as the incubation concentration of a sample required to bring the curve down to the point halfway between the top and bottom plateaus of the curve. Absolute IC50 here is defined as the incubation concentration of a sample that provokes 50% inhibition.

**Recombinant expression:** The Fab Sequences were sent to Sino Biological for protein expression. A human IgG1 Fc isotype was chosen for this purpose. The target gene was amplified by PCR and inserted into the appropriate expression vector for plasmid preparation. The sequence of the constructed vector was confirmed through sequencing. The correct plasmid was then amplified and passed on to downstream processes. The plasmids were mixed with transfection reagents in the optimal ratio for transient transfection and added to the flask or bioreactor containing HEK293 cells. The cells were cultured in a serum-free medium and maintained in Erlenmeyer Flasks on an orbital shaker or the bioreactor at 37°C for six days. The cell culture was centrifuged, and the supernatant was loaded onto an affinity purification protein G column at an appropriate flow rate. The purified proteins were analyzed by SDS-PAGE.

**Surface plasmon resonance, SPR:** Affinity analysis was performed by surface plasmon resonance (SPR) using the OpenSPR-XT instrument (Nicoya Lifesciences). RBD was immobilized in the active channel using the standard EDC/NHS amine coupling chemistry for amine coupling to high-capacity carboxyl sensors and 1 M Tris-based solution for reaction quenching (*SI Appendix* Fig. S2A). The optimized immobilization setup shown in *SI Appendix* Fig. S2A provided sufficient levels of RBD immobilization (3150 RU), well above the minimum requirement of the OpenSPR system.

PBS supplemented with 0.05% Tween20 was used as an immobilization and running buffer, showing minimal non-specific binding in the Reference Channel (*SI Appendix* Fig. S2B). A regeneration screen identified 10 mM Glycine-HCl pH 1.5 as a universal regeneration solution for subsequent analysis. Screening of a nanomolar concentration range for antibody analytes identified 100 nM as the optimal concentration for evaluating all candidates. Normalized sensorgrams for each concentration set were overlaid for comparative analysis (*SI Appendix* Fig. S2C), and measurement of complex stability was plotted as immediately post-injection (early stability) vs 30 sec after (late stability). The same experimental setup was used to perform an off-rate screen with 300 sec contact time and 600-sec dissociation of 100 nM analytes that were normalized for their relative binding responses and ranked based on relative dissociation percentages (*SI Appendix* Fig. S2D).

**Data analysis:**

***de Novo Protein Sequencing:*** Using Novor software (3), the MS/MS spectra in the bottom-up data were de novo sequenced. If a pair of de novo sequences overlap, and the overlapping region involves multiple (>1) amino acids that are mutations from germline sequences, the two sequences are regarded to be from the same antibody clone and merged to produce a longer sequence contig. This process is repeated until there are no more pairs available. The resulting contigs are aligned with the germline genes to discover candidate sequences covering each CDR. Note that some longer contigs may span more than one CDR. Examples of contig sequences retrieved from the assembly are in *SI Appendix* Table S3. MaxQuant software (Ver2.1.3) was used to calculate the relative quantity (peak area) of each unique peptide in the CDR contigs from each fraction in the separation experiments (NRT and native gel). For each separation experiment, a peptide’s normalized quantification vector was obtained by dividing its quantity in each fraction by the total quantity in all fractions, and the similarity score between contigs was calculated as the average Pearson correlation coefficient between every pair of unique peptides from the two contigs, respectively. The similarity scores from the two separation experiments were added up. A group of CDR contigs that have high total similarity scores to each other are clustered into one antibody clone. Finally, the CDR sequences from the same clone are aligned to the germline sequence, and other overlapping de novo peptides fill the gaps in the framework regions. The resulting fully covered heavy and light chains are reported as a candidate antibody clone. The similarity scores between CDR contigs and the MS coverage were used empirically to rank the candidate clones. The list of sequences can be found in Dataset S3.

***Nonreduced samples digestion:*** The identification of the S-S bridge containing peptides from nonreduced protease digestion was analyzed using pLink v2 (4). The software identification parameters were set as follows: Flow Type: Disulfide bond (HCD-SS); Enzyme: LysC-AspN; Peptide mass: 300-9000Da; Peptide length: 3-90; Fixed modification: Gln->pyro-Glu; and Variable modifications: Nethylmaleimide[C], Oxidation[M], Deamidated[N], Deamidated[Q], and Acetyl [ProteinN-term]. The protein database (Supplementary Information 3) contained the sequences of interest deduced from the polyclonal antibody analysis.

**References:**

1) Mann, M., Hendrickson, R.C., Pandey A., (2001) Analysis of proteins and proteomes by mass spectrometry, Annu Rev Biochem. 70, 437-73.
2) Krusemark, C., Frey, B., Smith, L. et al. (2011) Complete chemical modification of amine and acid functional groups of peptides, and small proteins Methods in Molecular Biology, 753, 77-91
3) B. Ma, Novor: Real-Time Peptide de Novo Sequencing Software. J Am Soc Mass Spectrom 26, 1885–1894 (2015).
4) Lu, S., Fan S. Yang B. et al. (2015) Mapping native disulfide bonds at a proteome scale Nature Methods12(4), 329-331
